# Adjuvants influence the maturation of VRC01-like antibodies during immunization

**DOI:** 10.1101/2022.06.10.495681

**Authors:** Maria L. Knudsen, Parul Agrawal, Anna MacCamy, K. Rachael Parks, Matthew D. Gray, Brittany N. Takushi, Arineh Khechaduri, Rhea N. Coler, Celia C. LaBranche, David Montefiori, Leonidas Stamatatos

**Affiliations:** Vaccine and Infectious Disease Division, Fred Hutchinson Cancer Research Center, Seattle, WA, 98109, USA; Department of Global Health, University of Washington, Seattle, 98195, WA, USA; Center for Global Infectious Disease Research, Seattle Children’s Research Institute, Seattle, WA; Department of Pediatrics, University of Washington School of Medicine, Seattle, WA; Division of Surgical Sciences, Duke University, Durham, NC, 27710, USA

## Abstract

Once naïve B cells expressing germline VRC01-class B cell receptors become activated by germline-targeting immunogens, they enter germinal centers and undergo affinity maturation. Booster immunizations with heterologous Envs are required for the full maturation of VRC01-class antibodies. Here, we examined whether and how three adjuvants, Poly(I:C), GLA-LSQ, or Rehydragel, that activate different pathways of the innate immune system, influence the rate and type of somatic mutations accumulated by VRC01-class BCRs that become activated by the germline-targeting 426c.Mod.Core immunogen and the heterologous HxB2.WT.Core booster immunogen. We report that although the adjuvant used had no influence on the durability of plasma antibody responses after the prime, it influenced the plasma VRC01 antibody titers after the boost and the accumulation of somatic mutations on the elicited VRC01 antibodies.

**ONE SENTENCE SUMMARY:** VRC01-class BCRs with different somatic mutations are being selected depending on the adjuvant used during immunization

## INTRODUCTION

Broadly neutralizing HIV-1 antibodies (bnAbs) have been isolated from HIV-1-infected individuals and the structures of diverse bnAbs, as well as those of their epitopes on the HIV-1 envelope glycoprotein trimeric spike (Env), have been well characterized (*1-3*). The epitopes targeted by bnAbs are located on multiple regions of Env, such as its apex (*4-9*), the CD4-binding site (CD4-BS) (*10-19*), the interface between the gp120 and gp41 subunits (*20-24*), the silent face of gp120 (*25, 26*), and the membrane proximal regions (MPER) of gp41 (along with lipid moieties into which MPER is embedded) (*22, 27-34*). In addition, some bnAbs recognize clusters of glycan moieties on the gp120 subunit (*35-37*), while other bnAbs recognize epitopes that contain both glycans and polypeptides (*38-48*).

bnAbs that recognize the same region of Env and share common genetic and structural features are grouped into ‘classes’ (*12*). The VRC01-class of antibodies recognize an epitope within the CD4-BS, and their heavy chains (HCs) are derived from the VH1-2*02 allele while their light chains (LCs) express 5 amino acid (5 aa) long CDRL3 domains (*14-16, 19, 49-53*). We note however that, not every Ab formed by the above-association of heavy and light chains targets the CD4-BS (*54*). They are among the most mutated bnAbs known (*55*) and can display up to 30 percent amino acid sequence divergence, yet they recognize their epitope on diverse Envs with similar angles of approach (*14-16*). VRC01-class bnAbs protect animals from experimental S/HIV infection (*56, 57*) and one mAb of that class, VRC01, was recently shown to prevent HIV-1 acquisition from susceptible, circulating primary HIV-1 viruses, in large phase 3 clinical trials (*58*). Thus, it is expected that VRC01-class bnAbs will be a component of the immune responses elicited by an effective HIV-1 vaccine.

Although VRC01-class bnAbs isolated from HIV-1 infected individuals bind diverse recombinant (rec) Envs and potently neutralize HIV-1 viruses from different clades, their unmutated forms do not (*15, 59-62*). So far, a natural Env (as expressed by a circulating virus) capable of binding the unmutated forms (‘germline’, gl) of VRC01-class antibodies has not been identified. It is believed that one reason for the lack of elicitation of VRC01-class antibodies through immunization with rec Envs is due to the failure of such proteins to activate naïve B cells that express the unmutated B cell receptor (BCR) precursors of VRC01-class antibodies (*60, 62, 63*).

We and others have designed Env-derived proteins capable of binding unmutated VRC01-class antibodies (*60, 62, 64-68*). Such constructs are commonly referred to as ‘germline-targeting’ (*63*). Germline-targeting proteins have been designed on the backbone of the outer domain of gp120 (*60, 64, 65*), the gp120 core (expressing both the inner and outer gp120 domains) (426c.Mod.Core) (*66, 68, 69*), and the entire extracellular portion of the viral Env (GT1.1) (*67*). A common feature of all such constructs is the elimination of the conserved N-linked glycosylation site at position N276 (Loop D of gp120), because glycans present at this position prevent glVRC01-antibodies/BCRs from binding to Env (*70*). However, additional obstacles on the HIV-1 Envs, including the lengths of the V1-V3 regions, prevent the engagement of germline VRC01-class BCRs by rec Envs (*66, 71*).

Orthologs of the human VH1-2*02 gene are not expressed in wild type animals such as mice, rats, rabbits, and non-human primates (*51*), thus the abilities of germline-targeting immunogens to activate naïve B cells expressing glVRC01-class BCRs have so far been evaluated in specifically engineered mice. Indeed, germline-targeting immunogens activate B cells expressing glVRC01-class antibodies *in vivo* (*65, 67, 68, 72-78*), although by themselves are not capable of guiding the proper maturation of VRC01-class BCRs towards their broadly neutralizing forms, through the accumulation of specific somatic mutations (*74, 75*). It is hypothesized that following the activation of naïve B cells expressing glVRC01-class BCRs by germline-targeting immunogens, sequential immunizations with diverse (and gradually more native-like rec Envs) will be necessary to achieve this goal (*74, 78*).

Adjuvants alter the magnitude as well as the quality of the immune response (*79-81*). However, the effect of adjuvant on the selection, expansion, and maturation of VRC01 B cell subclasses has not yet been investigated in detail, although there are some reports suggesting that different adjuvants may select for different HC/LC amino acids (*76*). As immunogens targeting the unmutated forms of VRC01-class antibodies enter clinical evaluation, identifying the adjuvant that promotes high levels of the appropriate somatic mutations will be important.

Here, we investigated how adjuvants affect the magnitude and duration of VRC01 antibody and B cell responses elicited by the 426c.Mod.Core germline-targeting immunogen (*66, 69*) and whether these responses can be boosted by a heterologous Env that does not recognize glVRC01-class BCRs on its own. To this end, we compared the antibody and B cell responses elicited by 426c.Mod.Core when expressed on Ferritin nanoparticles and adjuvanted with Polyinosinic-polycytidylic acid (Poly(I:C)), GLA-LSQ, or Rehydragel. Poly(I:C) is a double-stranded RNA analogue mimicking viral RNA that stimulates endosomal TLR3, GLA-LSQ consists of the TLR4 agonist glucopyranosyl lipid A (GLA) and the saponin QS21 in a lipid formulation, while Rehydragel, an aluminum hydroxide particulate formulation, does not activate known TLR pathways (*80*). We also investigated whether the adjuvants affect the maturation of VRC01-like B cells following a heterologous booster immunization with HxB2.WT.Core.

Our study indicates that long-lasting VRC01-like plasma antibody responses were generated irrespective of the adjuvant used during the prime immunization, but that the titers of plasma VRC01-like antibodies and the number of somatic mutations accumulated in VRC01-like antibodies were influenced by the adjuvant used during the heterologous boost immunization.

## RESULTS

### High titers of anti-CD4-binding site antibodies are elicited by a single immunization with 426c.Mod.Core irrespective of the adjuvant used

The mouse model we employed here is heterozygous for the human inferred glHC of the VRC01 mAb (VRC01^glHC^) (*76*). In this model, approximately 80% of B cells express the human transgene and all express mouse LCs (mLCs); ∼ 0.1% of which contain 5 aa-long CDRL3s. As a result, ∼ 0.08% of naïve B cells in these mice express VRC01-like BCRs, as compared to ∼0.01% in humans (*72, 76, 82*).

To test the effect of the adjuvant on the magnitude of the elicited antibody response, mice were immunized with 426c.Mod.Core Ferritin nanoparticles (24meric) adjuvanted with either Poly(I:C), GLA-LSQ, or Rehydragel. At the peak of the plasma antibody response (2 weeks post-immunization), all animals, irrespective of the adjuvant, generated autologous plasma antibodies, the majority of which targeted the CD4-BS on 426c.Mod.Core (autologous anti-CD4-BS antibodies) (**Fig. 1A**). At this point, the anti-426c.Mod.Core plasma antibody titers were significantly higher in the GLA-LSQ group than the Rehydragel group (p ≤ 0.05). However, no significant differences in the relative percentage of plasma antibodies against the CD4-BS of 426c.Mod.Core were observed at this time point. All animals, irrespective of the adjuvant used, also developed antibody responses against the heterologous germline-targeting immunogen, eOD-GT8 (*76*), at this early time point after immunization (**Fig. 1B**). The anti-eOD-GT8 plasma antibody responses exclusively targeted the VRC01 epitope on that protein, as they did not display reactivity to the version of the protein with a mutated VRC01 epitope (eOD-GT8 KO). We concluded that a single immunization with 426c.Mod.Core Ferritin nanoparticles, elicits high titers of autologous anti-CD4-BS antibodies, including antibodies that bind the VRC01 epitope, irrespective of the adjuvant used.

**Figure 1.**
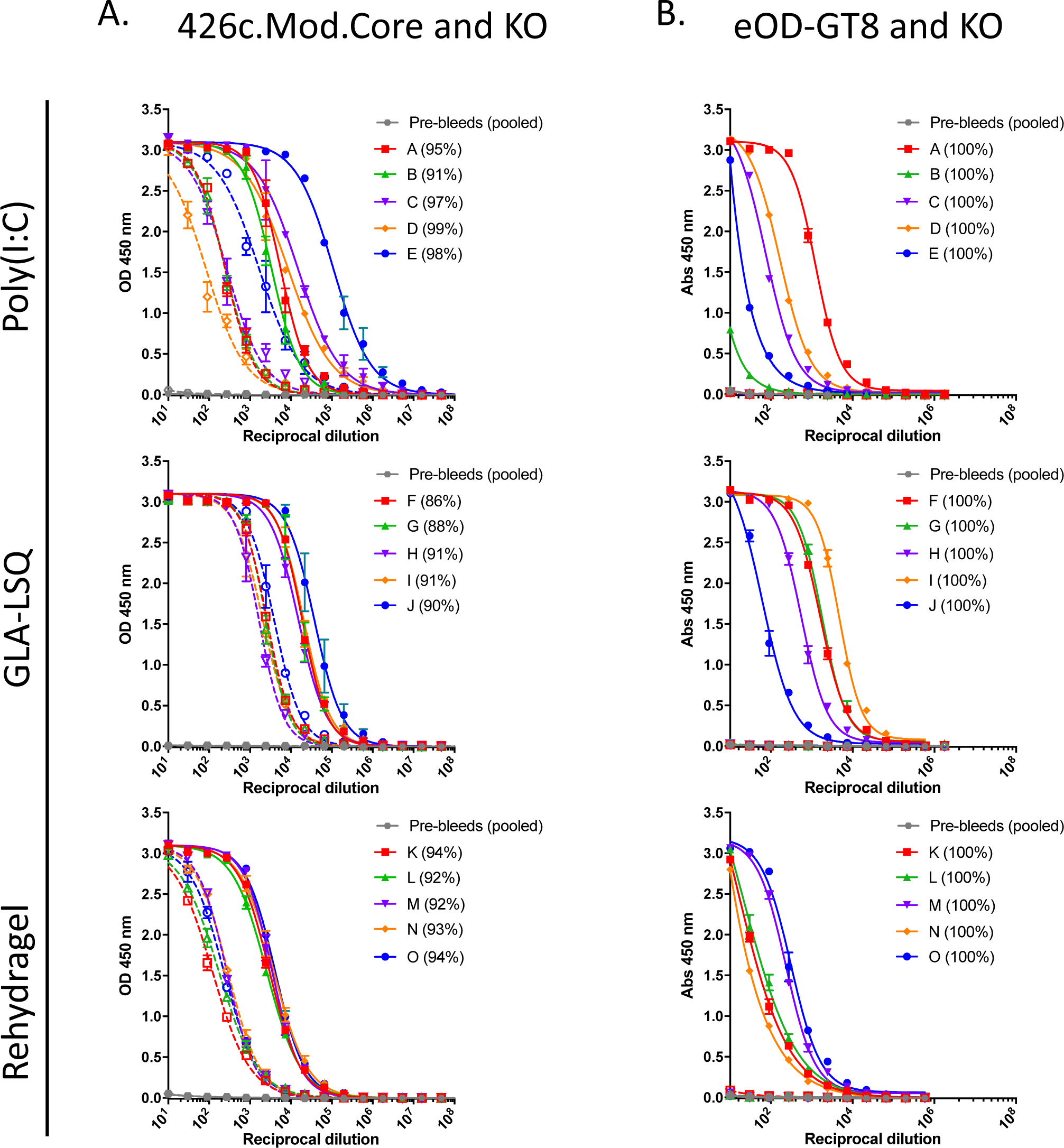
Ab responses at 2 weeks following 426c.Mod.Core Ferritin immunization. Mice (n = 5 per group) were immunized with 426c.Mod.Core Ferritin with either Poly(I:C), GLA-LSQ, or Rehydragel. Mice were bled at 2 weeks post-immunization and plasma was assayed by ELISA for binding against 426.Mod.Core (**A**) and eOD-GT8 (**B**) (full lines), as well as corresponding antigens with CD4-BS or VRC01 epitope knock-out (KO) (dotted lines). Figure legend indicates individual mouse, and percentage in parentheses indicates percent of response binding to CD4-BS. Pre-bleed samples from all animals (pool) was used as an internal control.

### Sustained autologous and heterologous HIV-1 Env antibody responses following a single immunization with 426c.Mod.Core

We next examined whether the longevity of the elicited antibody responses was affected by the adjuvant used. To this end, new groups of animals were immunized, and their responses were determined over a period of 22-23 weeks (**Fig. 2A**). The autologous plasma IgG titers were maintained at high levels over the course of observation (22-23 weeks). At this late timepoint post-immunization, the anti-426c.Mod.Core plasma antibody titers in the GLQ-LSQ and Poly(I:C) groups were not statistically different, although the antibody titers with Poly(I:C) were significantly higher than those in the Rehydragel group (p≤0.05).

**Figure 2.**
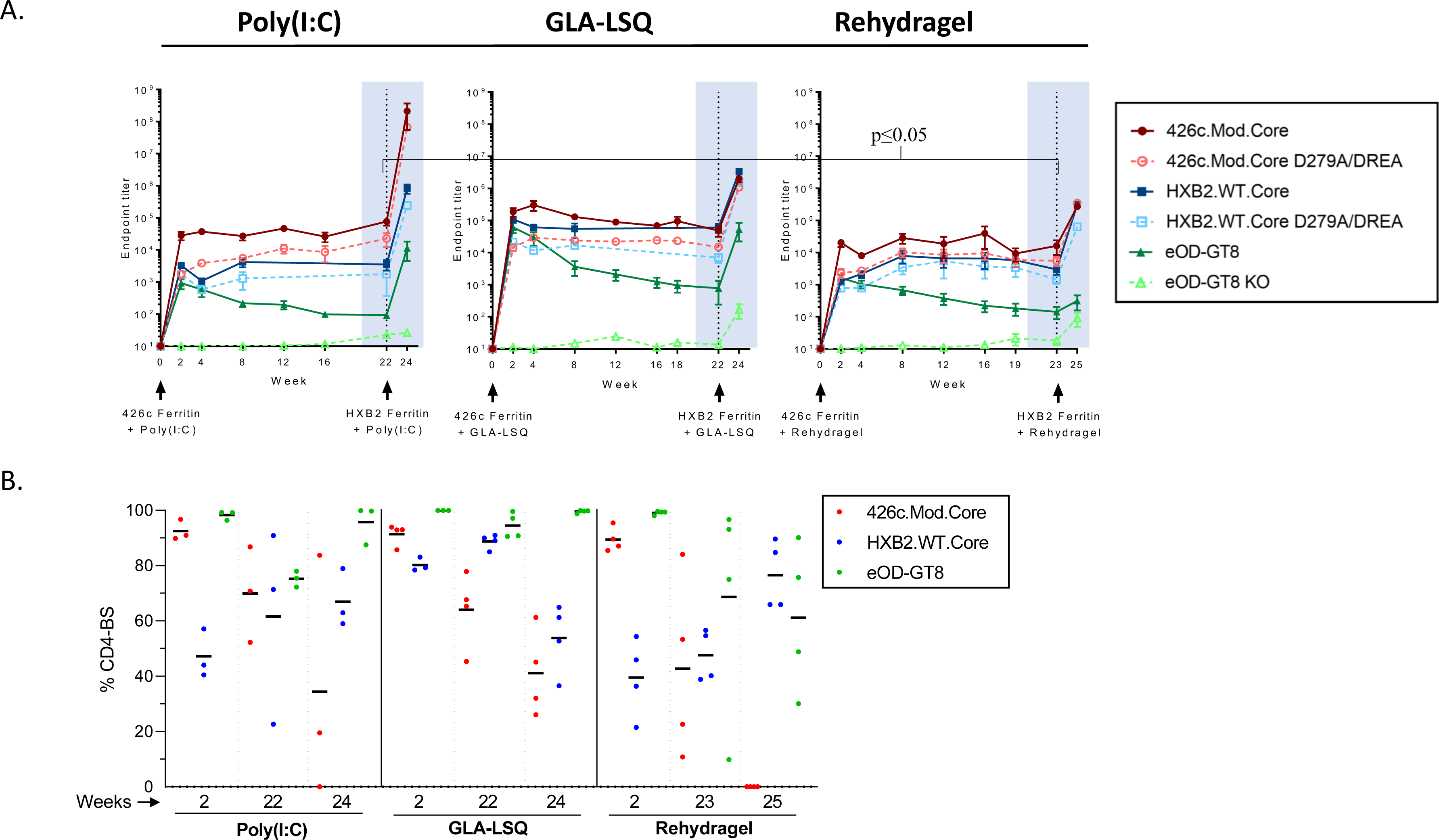
Ab responses following 426c.Mod.Core Ferritin prime and HXB2.WT.Core Ferritin boost immunization. Mice were primed with 426c.Mod.Core Ferritin at week 0 and boosted with HXB2.WT.Core Ferritin at week 22 (Poly(I:C) and GLA-LSQ groups), or week 23 (Rehydragel group). Mice were bled at the time points indicated in (A), and plasma was assayed for binding against the proteins listed in the legend by ELISA. Black dotted line indicates the time of the booster immunization. (**A**) endpoint titers against the indicated proteins over time. (**B**) percentage of anti-CD4-BS against 426c.Mod.Core (red), HxB2.WT.Core (blue) and VRC01 epitope on eOD-GT8 (green) at the indicated times with the indicated adjuvants. Each dot represents an animal.

We also examined whether the fraction of autologous anti-CD4-BS antibody responses remained constant over time. These responses peaked early following prime immunization (2 weeks post-immunization), but their relative proportions slowly decreased during the period of observation in all groups (**Fig. 2B**). At 22-23 weeks post-immunization, the autologous CD4-BS antibody responses represented ∼70% of the total autologous antibodies in the Poly(I:C) group (a drop of 24% from their peak value), 64% in the GLA-LSQ group (a drop of 30% from their peak value), and 43% in the Rehydragel group (a drop of 52% from their peak value) (**Fig. 2B**).

At the peak of the response, the anti-eOD-GT8 plasma antibody titers were 500-1,500-fold lower than the anti-426c.Mod.Core antibody titers and although the anti-eOD-GT8 titers gradually declined over time, they were always detectable (**Fig. 2A**). The majority of the anti-eOD-GT8 antibody responses targeted the VRC01 epitope on that protein for the duration of observation period as the reactivity to the eOD-GT8 KO was minimal (**Fig. 2A**).

The VRC01-like antibodies elicited by 426c.Mod.Core in this mouse model recognize heterologous, fully glycosylated wild type gp120 Core proteins (*68*). These constructs (termed ‘WT Cores’ for simplicity) are not recognized by glVRC01-class antibodies, but once activated by the 426c.Mod.Core, the VRC01-expressing B cells enter the germinal centers where their BCRs accumulate somatic mutations that allow them to bypass N276- and V5-associated glycans on some heterologous WT Core proteins. One of these WT Cores is the HxB2.WT.Core. Indeed, all animals generated durable anti-HxB2.WT.Core antibody responses following a single immunization with the 426c.Mod.Core (**Fig. 2A**), although the peak anti-HxB2.WT.Core responses were lower than those against 426c.Mod.Core. At 2 weeks post-immunization, the anti-HxB2.WT.Core antibody titers were ∼88%, ∼15% and 92% lower than the anti-426c.Mod.Core titers for Poly(I:C), GLA-LSQ and Rehydragel, respectively. The anti-HxB2.WT.Core plasma antibody titers remained stable for the duration of the observation, irrespective of the adjuvant used, and a fraction of these Ab target the CD4-BS. At week 2 post-immunization, the relative fraction of the plasma antibodies that recognized the CD4-BS on HxB2.WT.Core were ∼50% in the Poly(I:C) group, ∼ 80% in the GLA-LSQ group, and ∼40% in the Rehydragel group (**Fig. 2B**). The relative proportion of these heterologous anti-CD4-BS plasma antibodies remained unchanged over the duration of the observation. Thus, at 23 weeks post immunization, the relative fraction was ∼60% in the Poly(I:C) group, ∼90% in the GLA-LSQ group, and ∼50% in the Rehydragel group. We conclude that a single immunization with 426c.Mod.Core Ferritin nanoparticles elicits long-lasting autologous and heterologous anti-CD4-BS antibodies, a fraction of which target the VRC01 epitope.

### HXB2.WT.Core boosts autologous and heterologous anti-CD4-BS antibody responses primed by 426c.Mod.Core

We previously reported that immunization with a 7meric nanoparticle form of 426c.Mod.Core (426c.Mod.Core-C4b) adjuvanted with GLA-LSQ elicits VRC01-like antibody responses that are boosted by the heterologous HxB2.WT.Core (also in a 7meric nanoparticle form, HxB2.WT.Core-C4b), when administered 4 weeks after the prime immunogen, i.e., at the peak of the antibody response (*65, 68*). As a result, the VRC01-like antibodies isolated after this prime-boost immunization schema, have more somatic mutations and enhanced Env-binding affinities than the VRC01-like antibodies isolated after the 426c.Mod.Core prime immunization alone (*68*). Here, we examined how the anti-CD4-BS antibody responses, and more specifically the VRC01-like antibody responses elicited by the Ferritin nanoparticle form of 426c.Mod.Core, were affected by the adjuvant and by delaying the timing of the boost immunization with HxB2.WT.core Ferritin nanoparticles. To this end, animals were immunized with HxB2.WT.core Ferritin nanoparticles 22-23 weeks after the prime immunization.

This boost immunization resulted in increases in the plasma antibody responses against 426c.Mod.Core and HxB2.WT.Core in all three groups (**Fig. 2A**). The anti-426c.Mod.Core plasma antibody titers increased by ∼3,500 fold in the Poly(I:C) group, and by ∼1,500 fold in both the GLA-LSQ and the Rehydragel groups. The anti-HxB2.WT.Core titers increased by ∼3,000 fold in the Poly(I:C) group, by ∼2,000 fold in the GLA-LSQ group, and by ∼1,000 fold in the Rehydragel group. However, the relative proportions of anti-426c.Mod.Core and of anti-HxB2.WT.Core CD4-BS plasma antibodies were lower after the boost immunization than after the prime immunization (**Fig. 2B**). Two weeks following the heterologous boost immunization, the proportions of plasma antibodies targeting the 426c.Mod.Core CD4-BS were ∼34% in the Poly(I:C) group and ∼41% in the GLA-LSQ group, and no longer apparent in the Rehydragel group. The proportion of antibodies targeting the CD4-BS of the booster antigen HxB2.WT.Core, was ∼67% in the Poly(I:C) group, ∼54% in the GLA-LSQ group, and ∼77% in the Rehydragel group. Interestingly, the VRC01 plasma antibody titers, were boosted in the Poly(I:C) group by ∼2,000 fold and in the GLA-LSQ group by ∼1,500 fold, but not in the Rehydragel group. We concluded that Rehydragel may not be an optimal adjuvant to boost VRC01 plasma antibody responses elicited by the 426c.Mod.Core germline-targeting immunogen.

### The HXB2.WT.Core immunogen does not elicit VRC01-like plasma antibody responses

The HxB2.WT.Core does not bind glVRC01 antibodies (*68*), and it is thus expected that it will not activate naïve B cells expressing glVRC01-like BCRs. Hence, we hypothesized that the increase in VRC01-like plasma antibody responses observed during the booster immunization with HxB2.WT.Core in the case of the Poly(I:C) and GLA-LSQ adjuvants was not due to the activation of naïve VRC01-like B cells by the HxB2.WT.Core itself, but was due to a boosting of the memory VRC01-like B cells that transitioned to plasma cells secreting plasma antibodies at that stage. These memory VRC01-class B cells initially got activated by 426c.Mod.Core and their BCRs accumulated relevant somatic mutations that allowed them to bind HxB2.WT.Core.

To prove this point directly, we immunized mice with HxB2.WT.Core Ferritin nanoparticles adjuvanted with GLA-LSQ and examined the Env-recognition properties of the elicited plasma antibody. High autologous and anti-426c.Mod.Core plasma antibody titers were generated in all animals 2 weeks post immunization (**Supp Fig. 1A**). Between 20 and 70% of the anti-426c.Mod.Core, and between 60 and 80% of the anti-HxB2.WT.Core responses, targeted the corresponding CD4-BS (**Supp Fig. 1B**). Anti-eOD-GT8 plasma antibody responses were generated by 2 of 4 animals and were of lower magnitude than the anti-426c.Mod.Core or anti-HxB2.WT.Core plasma antibody responses (**Supp Fig. 1A**). However, the plasma antibody titers to eOD-GT8 and eOD-GT8 KO were either similar or the anti-eOD-GT8 KO titers were higher than the anti-eOD-GT8 titers (**Supp Fig. 1A**). These observations confirm that HxB2.WT.Core activates B cells that can target epitopes expressed on different Envs, some of which are located within the CD4-BS but is not capable of activating the B cells that produce VRC01 antibodies. This observation also supports the notion that HxB2.WT.Core can activate VRC01-like B cells only when they have accumulated relevant somatic mutations (*68*).

### Characterizing the impact of adjuvants on the maturation of VRC01-like antibodies

To prove directly that the boost immunization with HxB2.WT.Core led to the expansion of memory B cells expressing VRC01-like BCRs, we isolated such cells at 2 weeks after the prime immunization and two weeks after the boost and sequenced their VH/VL genes.

Two weeks after the prime immunization with 426c.Mod.Core Ferritin; 130, 206, and 172 class-switched Env+ B cells were individually isolated from the Poly(I:C), GLA-LSQ, and the Rehydragel group, respectively. 80 (62%), 59 (28%), and 87 (50%) of HCs were successfully sequenced from the Poly(I:C), GLA-LSQ, and Rehydragel groups; of which 95%, 97%, and 90%, were derived from VH1-2*02, respectively (**Fig. 3A**). The H35N mutation in CDRH2, that stabilizes the interaction between CDRH1 and CDRH3 on VRC01-class antibodies (*76*), was enriched by 21%, 81%, and 62%, of the VH1-2*02 HCs isolated from the Poly(I:C), GLA-LSQ, and Rehydragel groups, respectively (**Fig. 3B**). 41 (32%), 65 (32%), and 46 (27%) LCs, were successfully sequenced from the Poly(I:C), GLA-LSQ, and Rehydragel groups, respectively. 19 (46%) in the Poly(I:C) group, 52 (80%) in the GLA-LSQ group, and 31 (67%) in the Rehydragel group, contained the characteristic 5 aa-long CDRL3s (**Fig. 3C**). The majority of the 5 aa-long CDRL3s in all adjuvant groups were derived from the mouse 8-30*01 κV gene (**Fig. 3D**), as we previously reported (*68*). Antibodies expressing the VRC01 HC paired with other mouse LCs were also isolated (from the Poly(I:C) and Rehydragel groups) but only a few of these LCs expressed 5 aa-long CDRL3s (**Fig. 3D**). Within the 8-30*01 LCs, Glu96LC, a key CDRL3 feature of the mature VRC01-class antibodies, was detected in the Poly(I:C) group and more frequently in the GLA-LSQ group, but not in the Rehydragel group (**Fig. 3E,F**). These results suggest that adjuvants may differentially affect the selection of VRC01-like BCRs with particular amino acid mutations in both their HC and LCs.

**Figure 3.**
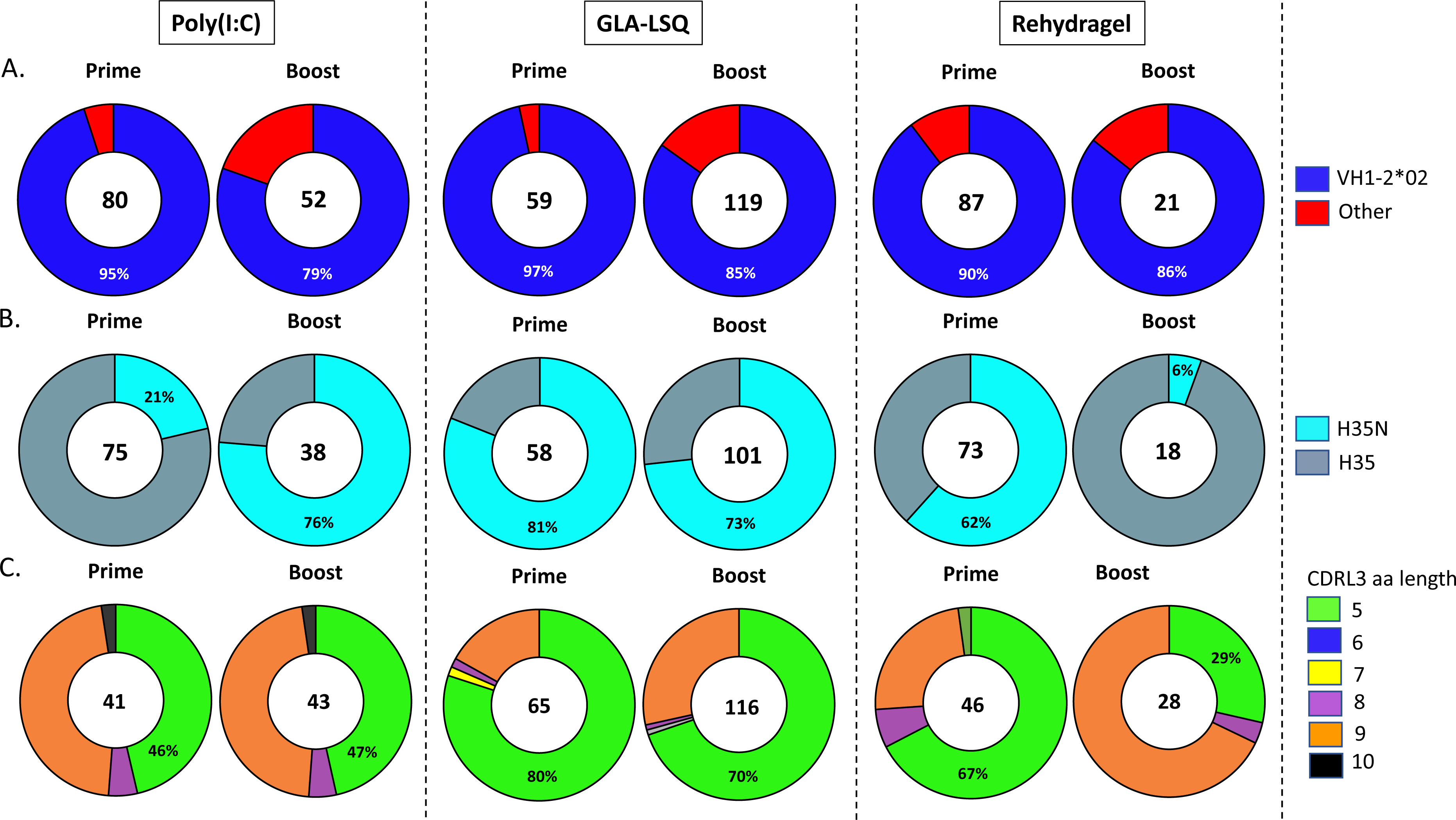
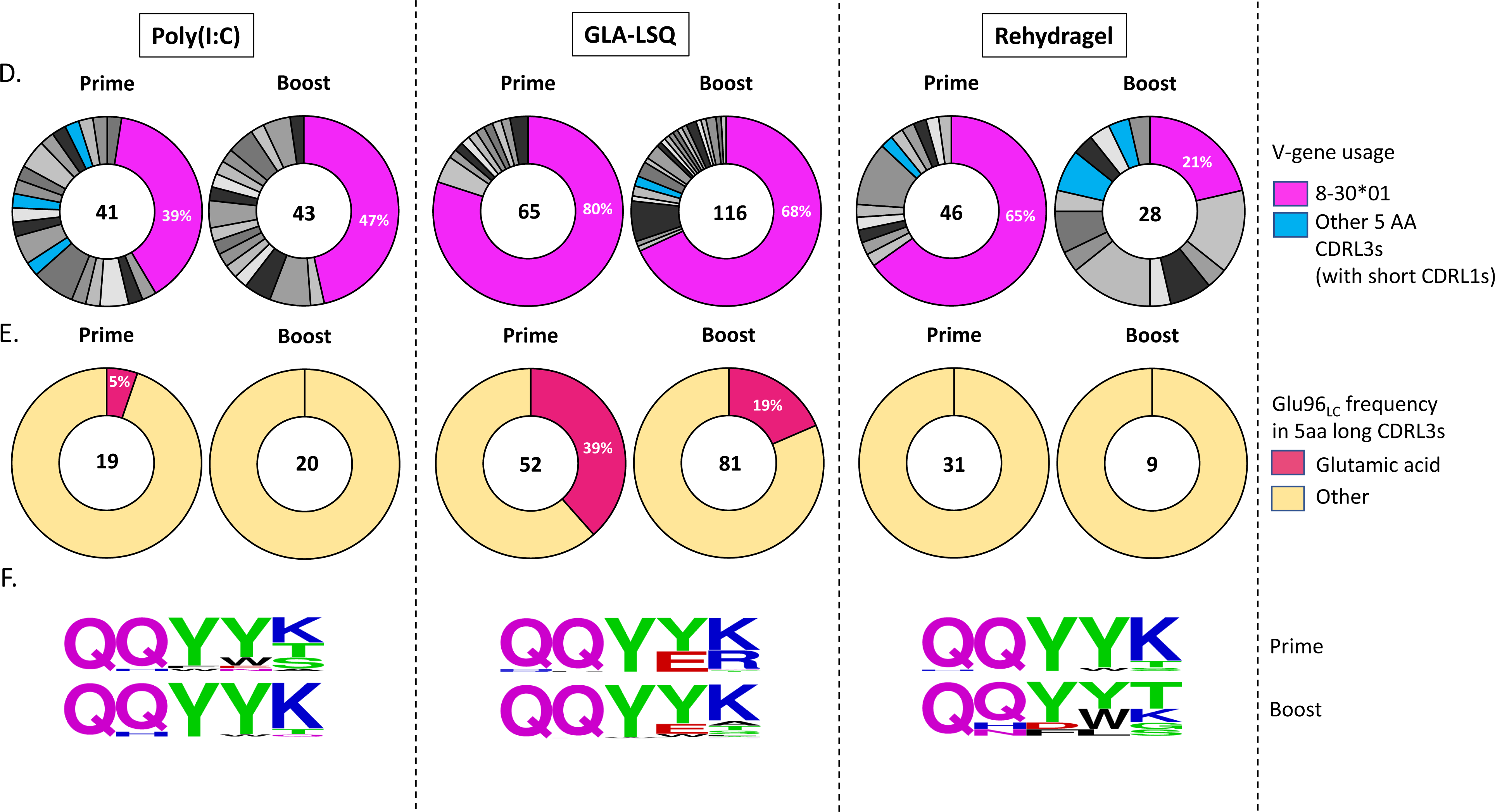
VH/VL sequence analysis from antibodies developed after the prime and boost immunizations. Pie charts indicate VH (**A,B**) and VL (**C-F**) gene usage from individually sorted B cells 2 weeks post immunization with 426c.Mod.Core (Prime) and 2 weeks post immunization with HxB2.WT.Core (Boost). The number of VH and VL sequences analyzed is shown in the middle of each pie chart. (**A**) VH-gene usage, (**B**) VHs with the H35N mutation. (**C**) aa length of the CDRL3 domains in the VL, (**D**) VL-gene usage. Shades of grey/black slices represent non 5 aa-long CDRL3s, k8-30*01 VLs are represented in pink while non k8-30*01 VLs with 5 aa long CDRL3s are indicated in blue, (**E**) Presence or absence of Glu96LC within the LC sequences with 5 aa-long CDRL3 domains, (**F**) Logo plots represent 5 aa-long CDRL3 sequences in the 8-30*01 VKs.

Two weeks following the boost immunization with HxB2.WT.Core, 124 B cells were isolated from the Poly(I:C) group, 248 B cells from the GLA-LSQ group, and 84 B cells from the Rehydragel group. 52 (42%) HCs and 43 (35%) LCs were successfully sequenced from the Poly(I:C) group, 119 (48%) HCs and 116 (47%) LCs from the GLA-LSQ group, and 21 (25%) HCs and 28 (33%) LCs from the Rehydragel group (**Fig. 3A,C**). Majority of the HC sequences (79% in the Poly(I:C) group, 85% in the GLA-LSQ group, and 86% in the Rehydragel group), expressed the VH1-2*02 gene (**Fig. 3A**). Interestingly the fraction of HCs with the H35N mutation in the Poly(I:C) group increased from 21% after the prime to 76% after the boost, while it decreased from 61% after the prime to ∼6% after the boost in the Rehydragel group and remained at similar levels in the GLA-LSQ group (**Fig. 3B**). 20 of 43 LCs (47%) in the Poly(I:C) group, 81 of 116 LCs (70%) in the GLA-LSQ group, and 8 of 28 (29%) LCs in the Rehydragel group, contained 5 aa-long CDRL3s (**Fig. 3C**). These frequencies were comparable to those observed after the prime immunization. As expected, the majority of the 5-aa CDRL3s were derived from the mouse 8-30*01 LC V gene (**Fig. 3D**). Noticeably, only the GLA-LSQ group enriched for 5-aa-CDRL3s containing Glu96LC after the boost (19%; **Fig. 3E,F**). These observations are in general agreement with those discussed above, in that the adjuvant influences which somatic mutations are selected by VRC01 BCRs.

After the prime immunization, a range of somatic mutations resulting in amino acid changes were observed in both the HCs and LCs (**Fig. 4**) in all three adjuvant groups. The mean numbers of HC amino acid mutations were: 2.3, 2 and 3.3 (**Fig. 4C**) and the mean numbers of LC amino acid mutations were: 1.9, 2.7 and 2.8 (**Fig. 4D**) for the Poly(I:C), GLA-LSQ, and Rehydragel groups respectively, following prime immunization. After the boost immunization, statistically significant increases in both nucleotide (**Fig. 4A**) and amino acid (**Fig. 4C**) mutations in the HCs were observed in the Poly(I:C) and GLA-LSQ groups (p≤0.001), but not in the Rehydragel group. Similarly, statistically significant increases in both nucleotide (**Fig. 4B**) and amino acid (**Fig. 4D**) mutations in the LCs were observed in the Poly(I:C) (p<0.01) and GLA-LSQ (p≤0.001) groups but not in the Rehydragel group. Additionally, the mean number of amino acid mutations in the three adjuvant groups differed significantly in the HCs (∼8.4, ∼6.6, and 2.8), and LCs (∼5.5, ∼5.4 and 2.2) for Poly(I:C), GLA-LSQ and Rehydragel, respectively. We concluded that Rehydragel may not be an optimal adjuvant for the maturation of VRC01 antibody responses.

**Figure 4.**
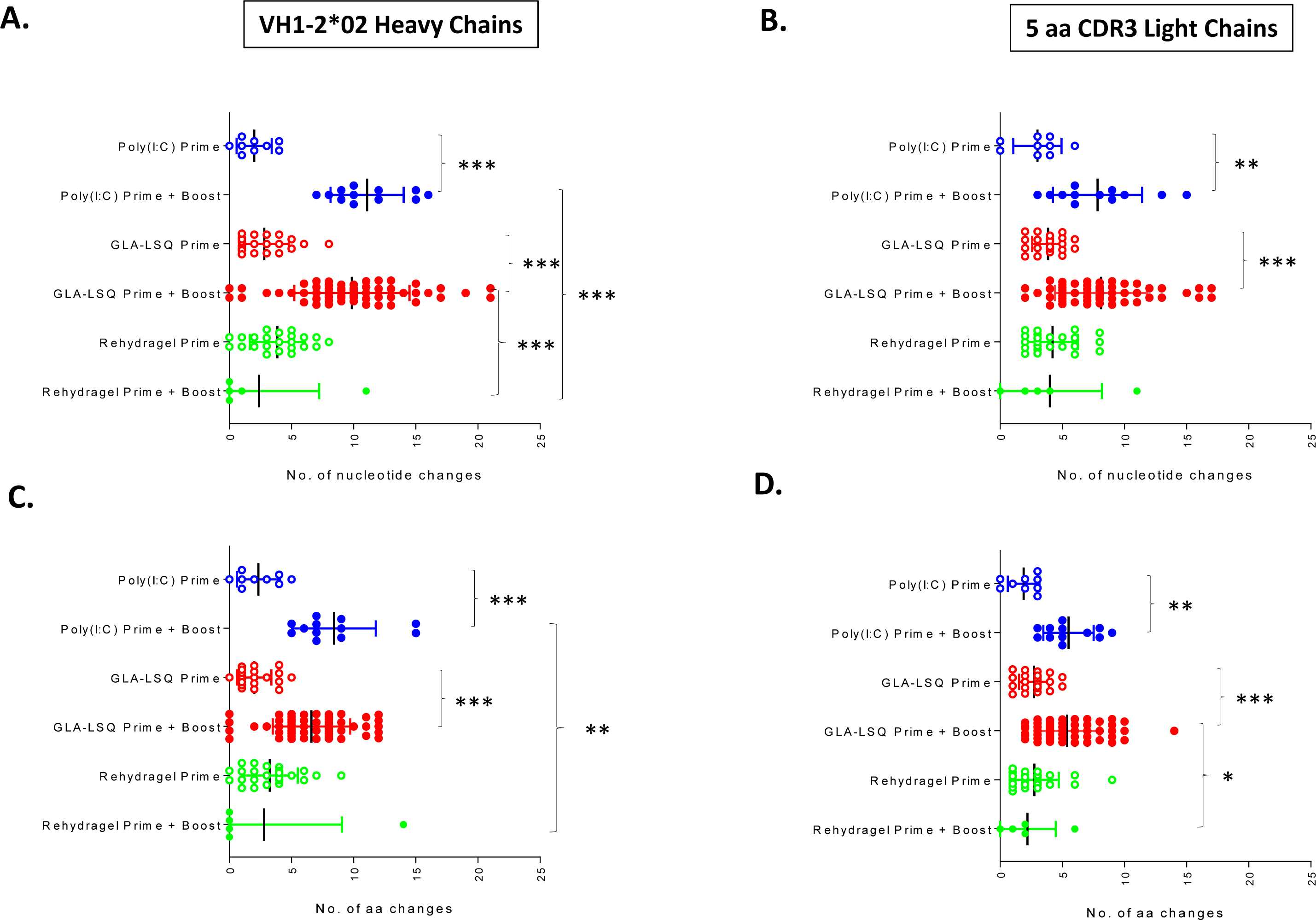
Number of nucleotide and amino acid changes in VRC01-like antibodies generated after the prime and after the boost immunizations. (**A**) number of nucleotide and (**B**) amino acid changes in the HC, and (**C**) number of nucleotide and (**D**) amino acid changes in the LC, of paired sequences isolated after the prime and boost immunizations with the indicated adjuvants. Significance was calculated using one-way ANOVA (Tukey’s multiple comparisons test). (*) ≤0.05; (**) ≤0.01 and (***) ≤0.001.

**Figure 5.**
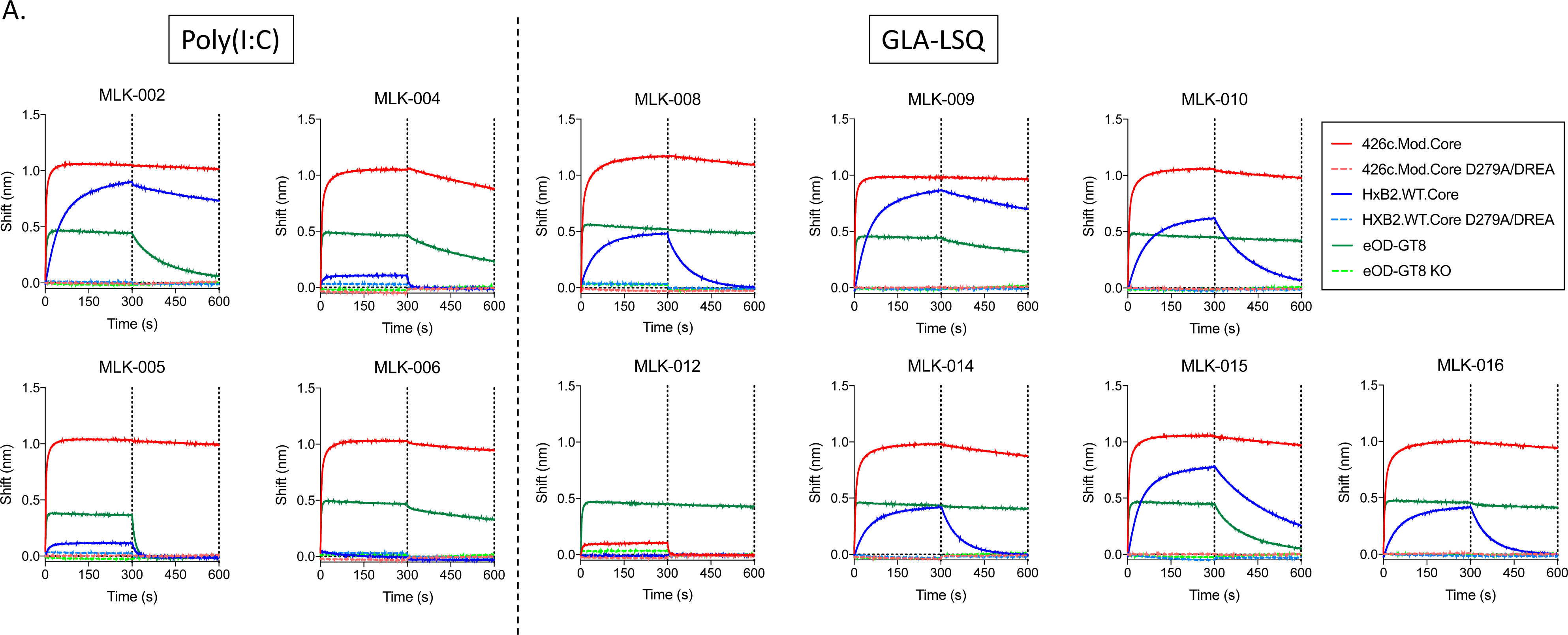
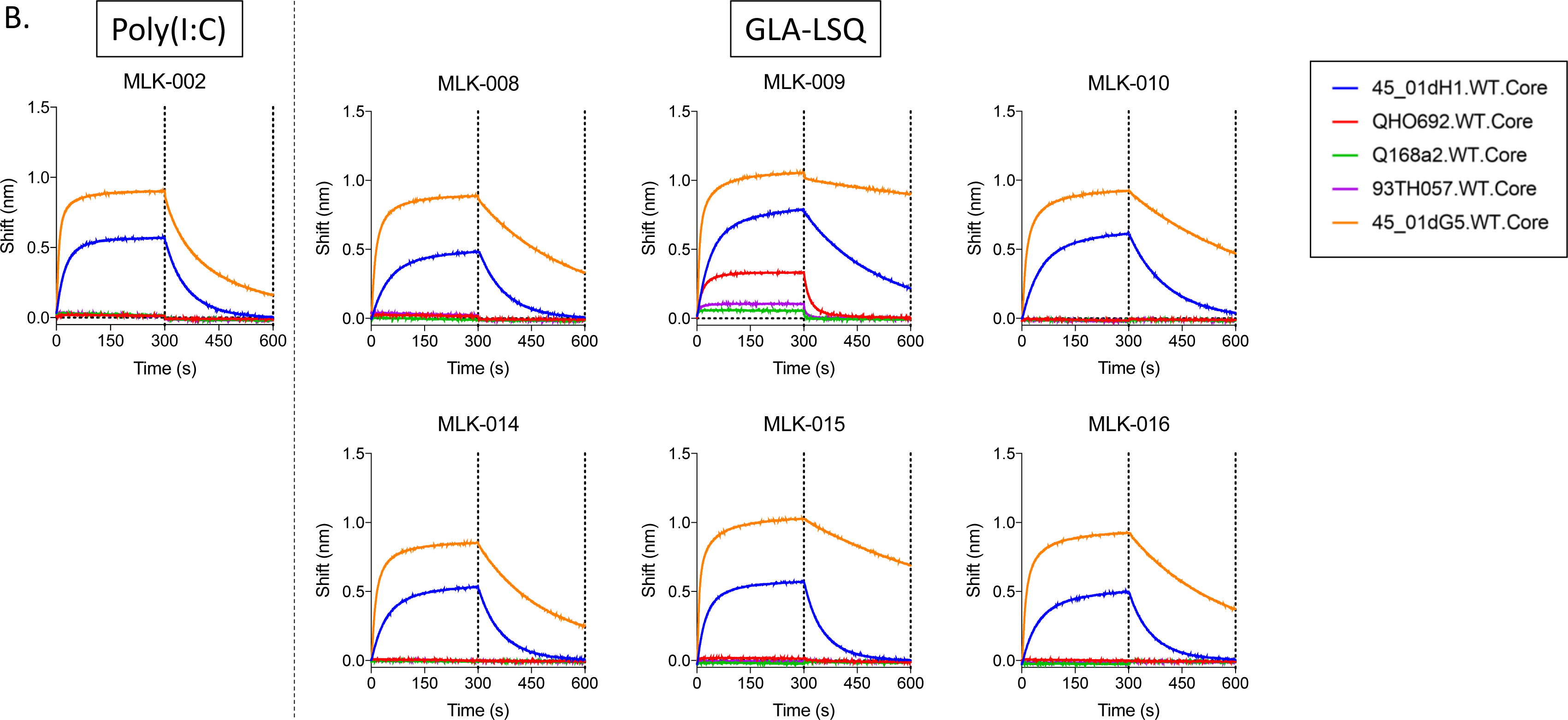
Binding properties of VRC01-like mAbs generated after the boost immunization. (**A**) Four VRC01-like mAb isolated from the Poly(I:C) group and seven VRC01-like mAbs isolated from the GLA-LSQ group were evaluated against the indicated soluble monomeric Envs. (**B**) those mAbs that displayed anti-HxB2.WT.Core reactivity were evaluated against the indicated five heterologous WT Cores. Information on these mAbs is provided in **Supplemental Figure 2**. Dotted lines indicate end of association and dissociation phases.

### Neutralizing properties of the plasma antibody responses after the boost immunization

Plasma IgG was purified 2 weeks following the boost immunization with HxB2.WT.Core Ferritin from all three adjuvant groups and was first tested against the tier 2, WT 426c virus produced either in 293T or GnTi-/-cells, but neither version was neutralized by any of the samples (**Fig. 6A**). We next tested these samples against a derivative of 426c that lacks 3 NLGS (N276, N460 and N463) (triple mutant, TM) expressed in GnTi-/- cells. The elimination of these NLGS renders the 426c susceptible to neutralization by some glVRC01-class antibodies (*62*).

**Figure 6.**
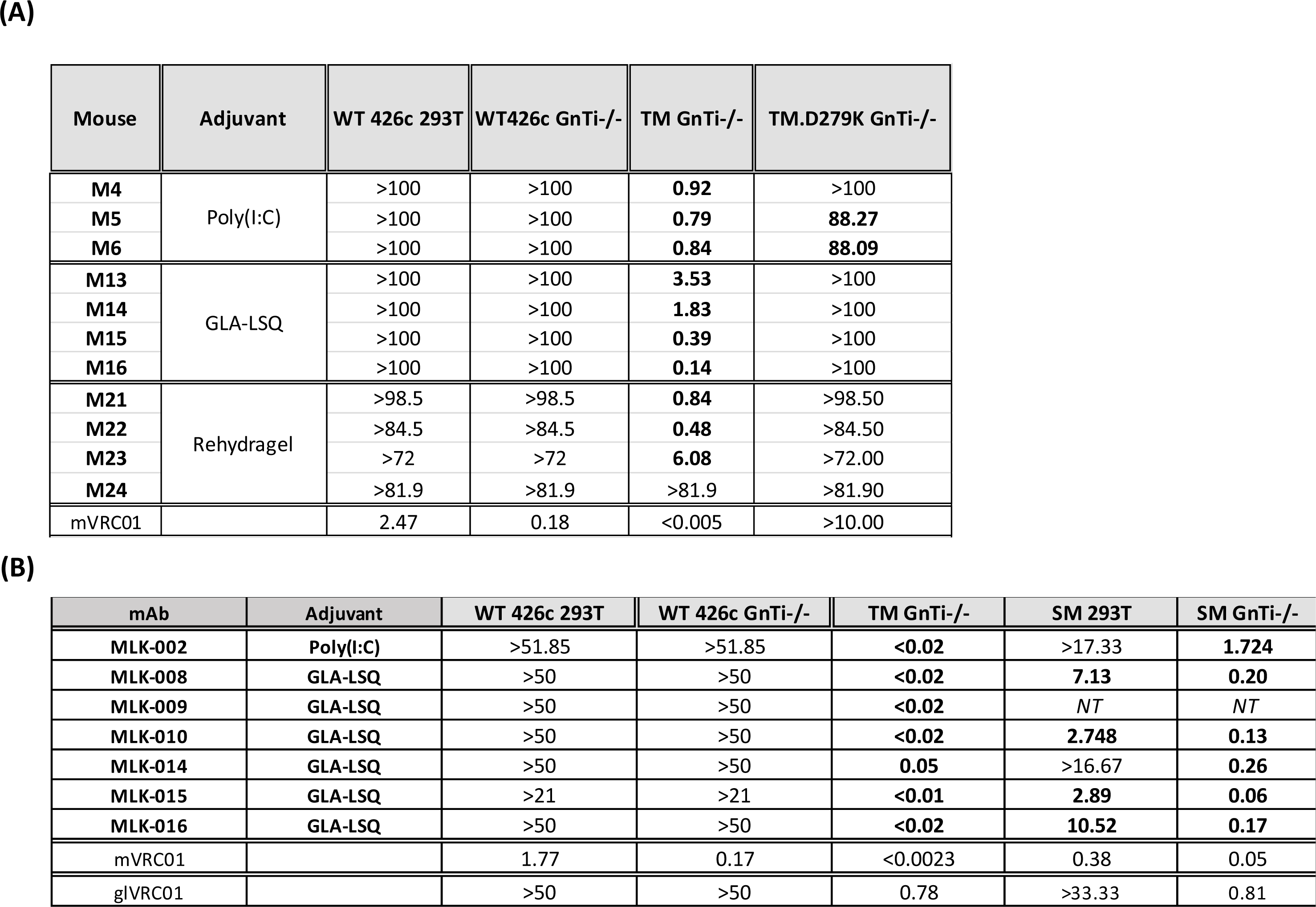
Neutralizing activities of plasma IgG and VRC01-like mAbs. (**A**) IgG was isolated from plasma collected 2 weeks following the booster immunization with HxB2.WT.Core from the indicated adjuvant groups and was evaluated for the presence of neutralizing antibodies against the WT 426c virus and the 426c virus whose Env lacks three NLGS (N276, N460 and N463), TM, and against its variant with the D279K mutation that abrogates the neutralizing activity of VRC01 antibodies. Viruses were expressed in 293 GnTi-/- cells. The WT 426c virus was also expressed in 293T cells. Values represent IC_50_ neutralization values in μg/ml. Neutralization IC50 values of these same viruses with the mature VRC01 mAb are included. Bold values indicate samples displaying neutralizing activity. (**B**) VRC01-like mAbs isolated after the boost immunization with HxB2.WT.Core from the Poly(I:C) (mAb MLK-002) and GLA-LSQ (all other mAbs) groups (**see** Figure 5 **for binding information**) were evaluated against the same viruses as the plasma samples as well as against the 426c variant whose Env only lacks the N276 NLGS (SM) expressed in 293T and 293 GnTi-/- cells. Values represent IC50 neutralization values in mg/ml. Neutralization IC50 values of these same viruses with the mature and germlineVRC01 mAbs are included. Bold values indicate neutralizing activity of VRC01 antibodies.

Ten of eleven samples neutralized this variant. Sample M24 that did not display anti-TM neutralizing activity was derived from the Rehydragel group. To confirm that the neutralizing activity was due to VRC01-like antibodies present in these samples, we tested the neutralizing activities of each sample against a derivative of TM that contains the D279K mutation, which abrogates the neutralizing activity of VRC01-class antibodies (*83*). In the GLA-LSQ or Rehydragel groups, the neutralizing activities were exclusively due to VRC01-like antibodies. In Poly(I:C) group, the neutralizing activity in one of the three samples (M4) was entirely due to VRC01-like antibodies, while in the remaining two samples (M5 and 6), the anti-TM neutralizing activity was primarily but not exclusively due to VRC01-like antibodies. We concluded that, irrespective of the adjuvant used, this prime-boost immunization schema elicits plasma VRC01-like antibody responses that cannot yet bypass the glycans present in Loop D (N276) and V5 (N460 and N463) on the WT 426c virus but can neutralize this virus when the three NLGS are absent.

### Neutralizing properties of monoclonal VRC01-like antibodies isolated after the boost immunization

To prove directly that VRC01-like antibodies were responsible for the neutralizing activities of plasma-derived polyclonal IgGs, we generated eleven VRC01-like mAbs from mice immunized in the presence of Poly(I:C) or GLA-LSQ, two weeks after the booster immunization with HxB2.WT.Core Ferritin (**Supp Fig. 2**). The mAbs express the human glVRC01HC paired with mouse k8-30*01 LCs expressing 5 aa-long CDRL3. With the exception of mAb MLK-012, they express amino acid mutations in both the HC and LCs. All mAbs recognized 426c.Mod.Core and eOD-GT8, but not their KO versions (**Fig. 5A**). With the exception of MLK-006 and MLK-012, all bound to HxB2.WT.Core but not its KO version. Among the mAbs that bound to HxB2.WT.Core, MLK-005 and MLK-006 displayed the weakest binding, while MLK-009 displayed the strongest binding.

The seven mAbs that bound to HxB2.WT.Core efficiently were also tested for binding to heterologous WT Cores (**Fig. 5B**). 45_01dG5 Env (clade B) is derived from a virus that circulated in patient 45, from which several VRC01-class antibodies have been isolated (including VRC01) (*84*). It naturally lacks the 276 NLGS and expresses one NLGS in its V5 region (position 470). All seven mAbs bound to this protein, with MLK-009 displaying the slowest off-rate compared to the remaining six mAbs. 45_01dH1 is derived from a virus circulating at a later time point in patient 45 and expresses the N276 NLGS, and, in addition to the N460 NLGS it expresses a second NLGS in V5 (position 476). Although all seven mAbs bound to that Env, they all displayed lower maximum binding and faster off rates than for 45_01dG5. Only mAb MLK-009 bound to the QH0692-derived WT core protein (clade B), and minimally bound to the Q168a2-derived WT Core protein (clade A) and 93TH057-derived WT Core protein (clade A/E). The remaining six mAbs did not bind to these proteins.

In agreement with the above discussed neutralization results obtained with polyclonal plasma IgGs, none of the seven VRC01-like mAbs neutralized the 426c WT virus, irrespective of whether it was produced in 293T or 293 GnTi-/- cells (**Fig. 6B**), but all neutralized the TM virus produced in 293 GnTi-/- cells. The mAbs also neutralized a 426c variant that only lacks the N276 NLGS (SM) when expressed in GnTi-/- cells and four of six mAbs neutralized this virus when expressed in 293T cells (mAbs MLK-002 and MLK-014 did not neutralize this virus).

Importantly, glVRC01 mAb did not neutralize this virus. The fact that all mAbs neutralized the TM virus but only 4 neutralized the SM virus, indicate that the glycans in V5 (N460 and N463) are significant obstacles for the maturing VRC01-like antibodies.

Overall, the data strongly suggest that the above immunization schema activates and initiates the maturation of VRC01-like B cells, but that the maturation process is incomplete as the elicited VRC01-like antibodies have not yet accumulated mutations that allow them to accommodate the glycans present on N276. We also note that these mAbs did not neutralize heterologous viruses whose N276 glycosylation site was eliminated by mutagenesis (N276Q) (data not shown). Thus, in addition to the N276-associated glycans, additional steric obstacles are present on heterologous Env that prevented the binding of these immature VRC01-like antibodies.

## DISCUSSION

The VRC01 antibody maturation process can be initiated by a single immunization with specifically designed Env-derived germline-targeting immunogens (*65, 68, 73, 76, 77, 85*). The completion of the maturation process, however, will require booster immunizations with distinct heterologous Envs (*74, 78*). In the transgenic animal model used here, that expresses the inferred human glHC of VRC01 paired with mLCs expressing 5 aa-long CDRL3 at a frequency of

∼0.08%, we and others reported on the incremental, but still incomplete, maturation of VRC01-like antibodies initiated by a prime immunization with a germline-targeting immunogen (426c.Mod.Core or eOD-GT8) followed by booster immunizations with 1-2 heterologous Env-derived proteins (*68, 73*). In a separate animal model, expressing both the inferred human glHC and human glLC of VRC01, Chen et al., reported on a more extensive maturation of VRC01-like antibodies using nine distinct immunogens administered sequentially that resulted in the isolation of VRC01-like antibodies displaying ∼50% neutralizing breadth (*74*). These observations support the overall ‘germline-targeting’ immunization approach (*86*) for the elicitation of VRC01 bnAbs and validate the potential of the utilized germline-targeting immunogens to initiate the antibody maturation process (*63*). It is however important to identify ways to optimize and accelerate this process. Here, we examined whether and how adjuvants may affect the maturation of VRC01-like antibody responses. The three adjuvants evaluated here were chosen because they activate different pathways of the innate response (Poly(I:C) via TLR3 (*87, 88*), GLA-LSQ via TLR4 (*89-91*), while Rehydragel acts independently of the TLR pathways (*92*).

Our results suggest that long-lived populations of plasma cells, including ones producing VRC01-like antibodies, can be generated by a single administration of the germline-targeting immunogen 426c.Mod.Core, irrespective of the adjuvant used. The fact that immunization with 426c.Mod.Core also elicited high and sustained plasma antibody responses against the heterologous HxB2.WT.Core suggests that this germline-targeting immunogen activates B cells that recognize conserved epitopes between these two proteins. A high fraction of these cross-reactive plasma antibodies targeted conserved elements of the CD4-BS. Interestingly the anti-CD4-BS plasma antibody titers (against 426c.Mod.Core or HxB2.WT.Core) decreased over time in all three adjuvant groups, while the total anti-426c.Mod.Core and anti-HxB2.WT.Core plasma antibody titers remained stable. This suggests that the numbers of plasma cells that produce anti-CD4-BS antibodies gradually decreased during the observation period while the numbers of plasma cells that target epitopes outside the CD4-BS gradually increased. A fraction of the anti-CD4-BS plasma antibody responses recognize the VRC01 epitope (expressed on eOD-GT8), and their titers also gradually decreased over the period of observation, in agreement with the overall decrease of the anti-CD4-BS plasma antibody responses.

While the plasma antibody responses to epitopes outside the CD4-BS increased by the boost immunization with HxB2.WT.Core, the anti-CD4-BS responses (to HxB2.WT.Core itself and to the prime immunogen 426c.Mod.Core) did not increase. This is not because the CD4-BS is not immunogenic on the HxB2.WT.Core. One possibility is that pre-existing anti-CD4-BS antibodies (elicited by the prime immunogen), bind the CD4-BS on the HxB2.WT.Core immunogen and prevent its recognition by naïve, or memory anti-CD4-BS B cells. That possibility however does not appear to apply to epitopes outside the CD4-BS. Importantly, however, in the case of Poly(I:C) and GLA-LSQ, an increase in plasma VRC01-like antibodies was observed after the boost immunization. This ‘boosting’ of VRC01-like plasma antibody responses suggests that an increase in the number of VRC01-like producing plasma cells took place by the heterologous immunization. The HxB2.WT.Core does not activate germline VRC01 B cells, and thus the observed increase in plasma VRC01 antibodies is due to the binding of HxB2.WT Core to B cells expressing VRC01-like BCRs that had been activated by the 426c.Mod.Core and had accumulated somatic mutations during the intervening period. The corresponding partially mutated plasma VRC01 antibodies represent a small fraction of the total antibodies in circulation at the time of the HxB2.WT.Core boost and thus do not prevent that protein from binding to VRC01-like B cells. Thus, despite a predominance of B cell responses to epitopes other than the VRC01 epitope on HxB2.WT.Core, boosting of VRC01-like B cells responses was possible in the presence of Poly(I:C) and GLA-LSQ, but not in the presence of Rehydragel. Presently, we do not know why the plasma VRC01-like B cell response were not boosted in the Rehydragel group, but we speculate that is related to the activation of different innate pathways.

Importantly, the adjuvant used affected the rate at which activated VRC01-class B cells accumulated somatic mutations in their VH/VL genes. Significantly more nucleotide and amino acid mutations were present in the VRC01-like antibodies isolated after the boost from animals that received immunogens adjuvanted with Poly(I:C) or GLA-LSQ than Rehydragel. It is possible that adjuvants affect the activation of CD4+ T cells that assist B cells in GCs, and/or the adjuvants directly affect these B cells. We expect the above effect to not be specific for VRC01 B cells, but to all B cells that became activated by the two immunogens used here.

As a result of the different rates of somatic mutations observed in the Rehydragel group and the GLA-LSQ or Ploy(I:C) groups, B cells expressing VRC01-like BCRs with particular amino acids at key positions were selected. Thus, the Glu96LC amino acid was present in VRC01-like B cells found in the Poly(I:C) and GLA-LSQ groups following the prime immunization with 426c.Mod.Core Ferritin, but not in the Rehydragel group. Glu96LC is found in all presently known human VRC01-class bnAbs and is the product of affinity maturation (*51*). It forms hydrogen bonds with the side chain of Asn280 in Loop D and with the backbone amide of Gly459 in V5 and contributes to increased antibody-Env affinity (*51, 70*). Glu96LC was only present in VRC01-like antibodies generated from the GLA-LSQ group after the boost immunization with HxB2.WT.Core Ferritin. Similarly, a higher frequency of VRC01-like antibodies isolated after the boost immunization expressed N35HC (rather than H) in the case of Poly(I:C) and GLA-LSQ than Rehydragel. The H35N mutation allows for a hydrogen bond formation with N100a in CDRH3, thus improving the HC/LC interaction (*76*).

In sum, our study provides direct evidence that adjuvants influence the activation and maturation of B cells expressing VRC01-like BCRs. As the elicitation of fully matured VRC01-like antibodies requires the accumulation of an extensive number of mutations in both the VH and VL genes (*55*) and several immunogens have to be administered in a particular sequence to select VRC01 BCRs with particular mutations at each immunization step (*74*), our study highlights the importance that adjuvants have in the selection of the appropriate mutations.

## AUTHOR CONTRIBUTIONS

Conceptualization: LS, MLK, PA

Methodology: LS, MLK, PA

Validation: LS, MLK, PA

Formal analysis: MLK, PA, DM

Investigation: MLK, PA, AM, K.R.P, M.D.G, B.N.T, AK, RNC

Writing – Original Draft: LS, MLK, PA

Writing – Review Editing: All Visualization: MLK, PA

Supervision: LS

Project Administration: LS Funding Acquisition: LS

## DATA AND MATERIAL AVILABILITY

All materials and reagents will be available. In certain cases, appropriate MTAs will need to be signed.

## SUPPLEMENT MATERIAL

### MATERIALS and METHODS

#### Recombinant HIV-1 Envelope proteins

Recombinant HIV-1 envelope proteins (rec Envs) were expressed by transient transfection in HEK 293-F cells and then purified directly from conditioned media as we previously described (*66*). The “CD4-BS knockout” (KO) versions of rec Envs contain the D279A, D368R and E370A mutations (D279A/DREA). In the case of eOD-GT8, the KO version contains the D368R mutation and the amino acids DWRD at positions 276-279 were substituted by NFTA. Purified Env proteins were aliquoted in PBS and stored frozen in −20°C until further use. Ferritin nanoparticles expressing 426c.Mod.Core and HxB2.WT.Core were produced and purified as previously described (*68*). They were stored at 4°C. Tetramers of Avi-tagged eOD-GT8 and of eOD-GT8 KO were generated as previously reported (*68*).

#### Adjuvants

Polyinosinic-polycytidylic acid (Poly(I:C)) was obtained from InvivoGene. GLA-LSQ was provided by the Infectious Disease Research Institute (IDRI) and liposomal formulations of GLA were prepared as previously published (*93*). Rehydragel was provided by the NIAID.

#### Mice and Immunizations

Knock-in mice expressing the inferred germline HC of the human VRC01 Ab (VRC01^glH^) and endogenous mouse LCs (*76*) were bred and kept at the Fred Hutchinson Cancer Research Center. Mice were 6-12 weeks old at the initiation of experiments. Proteins and adjuvants were diluted in PBS and administered intramuscularly with 50 μl in each hind leg in the gastrocnemius muscle (total volume 100 μl/mouse). Env antigens were administered at 50 (GLA-LSQ groups) or 60 μg (Poly(I:C) and Rehydragel groups), and adjuvants at 6 0 μg for Poly(I:C), 50 μl GLA-LSQ (containing 5 μg TLR4 agonist and 2 μg Saponin for GLA-LSQ), and 100 μg for Rehydragel. Blood was collected by the retroorbital route into tubes containing 25 μl citrate-phosphate- dextrose solution (Sigma-Aldrich). Terminal bleeds were collected by cardiac puncture into tubes containing 100 μl citrate-phosphate-dextrose solution. Organs were harvested into cold IMDM media (Gibco).

#### Organ processing

Spleens and lymph nodes were first mashed through 70-μm-pore-size nylon cell strainers (Falcon) to obtain a single cell suspension. Cells from lymph nodes were then washed twice with PBS, while splenocytes were first treated with Red Blood Cell Lysing Buffer Hybri-Max (Sigma-Aldrich) for 1.5-2 min followed by two washes with PBS. Cells were resuspended in FBS supplemented with 10% DMSO and frozen in a Mr. Frosty Freezing Container (Thermo Fisher Scientific) at -80°C overnight, then moved to liquid nitrogen for storage until use.

#### Single B cell sorting

Splenocytes or lymph node cells were thawed and first stained with Fc-block (2.4G2; BD Biosciences), 1 pmol Gb phycoerythrin (PE)-DyLight (DL)650 (PE-Decoy) and 3 pmol Gb allophycocyanin (APC)-DL755 (APC-Decoy) diluted in fluorescence-activated cell sorting (FACS) buffer (2% FBS, 1 mM EDTA in PBS). PE-Decoy and APC-Decoy were made by first combining streptavidin-PE (SA-PE) or streptavidin APC (SA-APC) (Prozyme) with a DL NHS ester (650 or 755; Thermo Fisher Scientific). 1 pmol eOD-GT8-PE tetramer and 3 pmol eOD- GT8 KO-APC tetramers were added to the cells and incubated on ice for 25 mins. Samples were then washed and incubated with anti-PE and anti-APC MicroBeads (both Miltenyi Biotec) after which decoy- and tetramer-positive cells were enriched by putting the samples through magnetic LS columns positioned on a QuadroMACS separator (all from Miltenyi Biotec) (19) Non-bound cells present in the flow-through were collected to use as controls. After a wash, samples (both bound and non-bound fractions) were incubated with Fixable Viability Dye eFluor 506 (eBioscience, ThermoFisher Scientific) and the following Abs diluted in Brilliant Stain Buffer (BD Biosciences): Anti-IgG1-fluorescein isothiocyanate (FITC; A85-1), anti-IgG2b-FITC (R12- 3; both from BD Biosciences), anti-IgG2c-FITC (polyclonal; Bio-Rad), anti-IgG3-FITC (R40- 82; BD Biosciences), anti-IgD-PerCP-Cy5.5 (11-26c.2a; BioLegend), anti-CD3-Brilliant Violet (BV)510 (145-2C11), anti-CD4-BV510 (RM4-5), anti-Gr-1-BV510 (RB6-8C5; all from BD Biosciences), anti-F4/80-BV510 (BM8), anti-IgM-BV605 (RMM-1; both from BioLegend), anti- CD19-BV650 (1D3), anti-B220-BV786 (RA3-6B2; both BD Biosciences). Samples were washed and resuspended in FACS buffer, AccuCheck Counting Beads (Thermo Fisher Scientific) were added, and the samples were then evaluated on a FACSAria II (BD Biosciences). Stained UltraComp eBeads (Thermo Fisher Scientific) were used for compensating voltages. Non-bound cells collected in the enrichment step described above were used for setting up gates. Samples were single cell-sorted into 96-well skirted plates (Eppendorf) and stored at - 80°C until further processing. 20 μl lysis buffer per well (20 U Rnase out, SSIV buffer, 6.25 μM DTT (all from Thermo Fisher Scientific), 0.3% Igepal) was added either immediately before sorting cells, or after thawing plates for further processing.

#### VH/VL gene sequencing

Amplification and sequencing of the antibody VH/VL genes was performed as we previously described (*65, 68, 76*). Briefly, cDNA was generated by adding 6 μl per well of a mix containing Random Primers, 3.3 mM dNTPs, 200 U Superscript IV Reverse Transcriptase (all from Thermo Fisher Scientific) and with PCR program 42°C 10 min, 25°C 10 min, 50°C 60 min, 94°C 5 min. cDNA was amplified and sequenced by two rounds of PCR using primers listed in **Supplemental Table 1** using PCR program: 94°C 5 min, 50 × (94°C 30 sec; X°C for 30 sec (X being the corresponding annealing temperature of the primers) and 72°C for 55 sec), and 72°C 10 min. All reactions were performed in a 40 μl volume with 2.4 U HotStarTaq Plus DNA Polymerase (Qiagen), 0.24 μM of a 5’ primers pool, 0.24 μM of a 3’ primer pool (all from Integrated DNA Technologies), 0.35 mM dNTPs. Aliquots from each well were subjected to agarose gel electrophoresis, which were then treated with ExoSAP-IT PCR Product Cleanup Reagent (Applied Biosystems, Thermo Fisher Scientific) and sequenced by Sanger sequencing using the primers indicated in **Supplemental Table 1**, as previously reported (*76, 94*).VH and VL sequences were analyzed using the Geneious software (Biomatters, Ltd.) and the online IMGT/V-QUEST tool (*95*). To calculate numbers of nucleotide and amino acid mutations, sequenced HC and LC pairs were aligned against the VH regions of the VRC01 gH knocked-in sequence and LC reference sequences obtained from IMGT/V-QUEST, respectively, using sequences starting from CDR1 to CDR3.

#### VH/VL cloning and antibody expression

DNA products from the 1^st^ round of PCR were used as templates for gene-specific PCR to amplify the gene of interest and add ligation sites to allow for insertion of the DNA fragment into the human IgG1 vectors: ptt3 for κ light chain (*96*) and PMN 4-341 for γ heavy chain (23). Each gene-specific PCR reaction consisted of 0.5 μl of each 10 μM 5’ and 3’ primer, 22.5 μl Accuprime Pfx Supermix (Thermo Fisher Scientific) and 1.5 μl of 1^st^ round PCR product. The gene-specific PCR product was then infused into cut IgG1 vector in a 2.5 μl volume reaction containing 12.5 ng of cut vector, 50 ng of insert, 0.5 μl of 5× Infusion enzyme (Takara Bio).

Competent *E. coli* cells were transformed with the entire reaction and plated onto ampicillin agar plates. Colonies were picked and grown in LB broth containing ampicillin, and DNA was extracted and purified using QIAprep Spin Miniprep Kit (Qiagen). 293E cells were then transfected with equal amounts of HC and LC DNA as well as 293F transfection reagent (Millipore Sigma) and grown for 5-7 days, at which time Abs were purified from cell supernatants using Pierce Protein A agarose beads (Thermo Fisher Scientific). Abs were eluted with 0.1 M Citric Acid into 1 M Tris buffer followed by buffer exchange into PBS using an Amicon centrifugal filter (Millipore Sigma).

#### Purification of IgG from mouse plasma

IgGs were purified from mouse plasma using Protein G HP-Ab Spin Trap columns (GE Healthcare Life Sciences) according to the manufacturer’s protocol. Eluted antibodies were buffer-exchanged into PBS using Amicon Ultra-4 centrifugal filter units (30K, Merck Millipore Ltd.).

#### ELISA

384 well ELISA plates (Thermo Fisher Scientific) were coated with 0.1 μM his/avi-tagged protein (426c.Mod.Core, 426c.Mod.Core KO, HXB2.WT.Core, HXB2.WT.Core KO, eOD-GT8, eOD-GT8-KO) diluted in 0.1 M sodium bicarbonate, at room temperature (RT) overnight. Plates were then washed four times with wash buffer (PBS plus 0.02% Tween20) using a microplate washer (BioTek) and incubated with block buffer (10% milk, 0.03% Tween20 in PBS) for 1-2h at 37°C. Plates were washed, mouse plasma added, and serially diluted (1:3) in block buffer.

After 1h incubation at 37°C, plates were washed, and horse radish peroxidase-conjugated goat anti-mouse IgG (BioLegend) was added and incubated for 1h at 37°C. After a final wash, SureBlue Reserve TMB Microwell Peroxidase Substrate (KPL Inc.) was added to the plates for 5 mins. The reaction was stopped with 1 N H2SO4, and the optical density (OD) was read at 450 nm with a SpectraMax M2 Microplate reader (Molecular Devices). The average OD of blank wells from the same plate were subtracted from all wells before analysis.

#### Biolayer interferometry

Biolayer interferometry (BLI) assays were performed on the Octet Red instrument (ForteBio) as previously described (*65, 68*). Briefly, anti-human IgG FC capture biosensors (ForteBio) were used to immobilize mAbs (20 μg/μl in PBS) for 5 min, followed by baseline interference reading for 60 s in kinetics buffer (PBS, 0.01% BSA, 0.02% Tween-20, 0.005% NaN3). Sensors were then immersed into wells containing Env Core monomers (2 μM) diluted in kinetics buffer for 300 s (association phase) and another 300 s (dissociation phase). All measurements were corrected by subtracting the signal obtained from simultaneous tracing of the corresponding Env using an irrelevant IgG Abs in place of the mAbs tested. Curve fitting was performed using the Data analysis software (ForteBio).

#### TZM-bl neutralization assay

Plasma IgG and individual mAbs were tested for neutralization against a panel of selected HIV-1 pseudoviruses using TZM-bl target cells, as previously described (*97*). Germline and mature VRC01 mAbs were used as controls in every assay.

#### Statistics

For pairwise comparisons between two groups, the unpaired *t*-test was used. For comparisons between three or more groups, the ANOVA test was used for *a priori* analyses, followed by Tukey’s or Sidak’s multiple comparison’s test for post hoc analyses. A *P* value of ≤ 0.05 was considered statistically significant. Statistical analyses were carried out using the GraphPad Prism software (GraphPad Software).

### SUPPLEMENTARY FIGURE LEGENDS

**Supplementary Figure 1.**
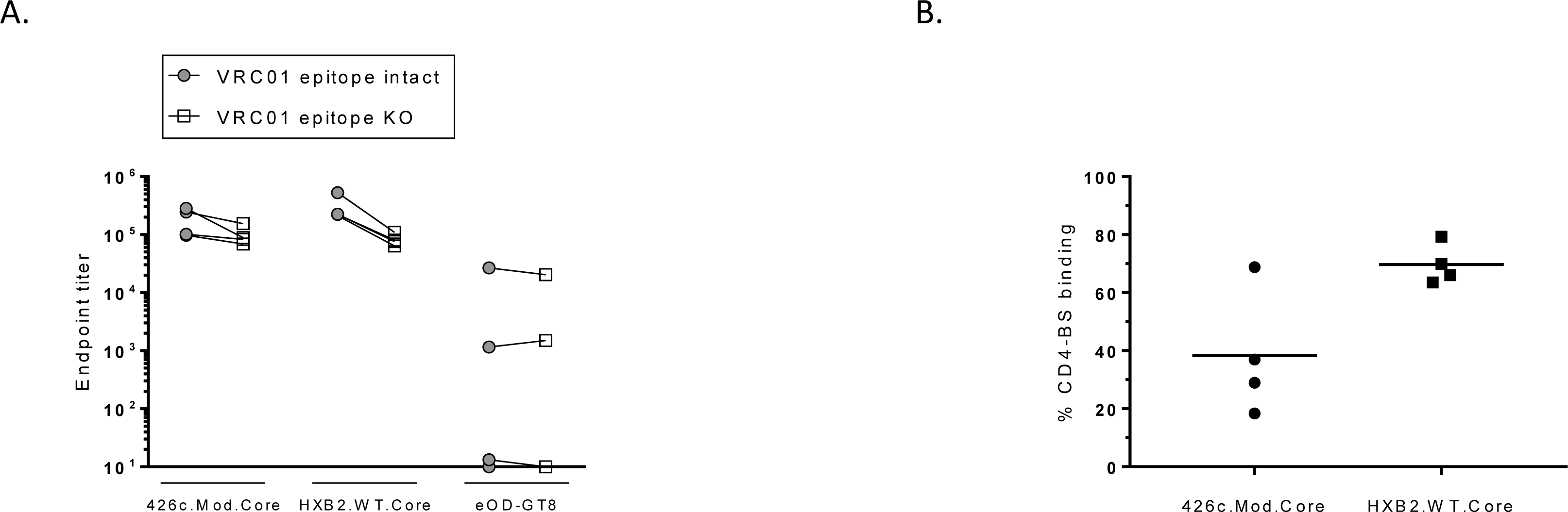
Plasma antibody responses elicited by HXB2.WT.Core. (**A**) Mice were immunized with HXB2.WT.Core Ferritin adjuvanted with GLA-LSQ, and the plasma antibody responses were evaluated 2 weeks later against the indicated Env proteins (closed symbols) and their corresponding CD4-BS KO versions (open symbols). Each symbol represents one mouse, and lines connect data from the same mouse. (**B**) Percentages of plasma antibodies binding to the CD4-BS of the indicated proteins (derived from the results in panel (**A**)). Horizontal lines represent mean values of the indicated groups.

**Supplementary Figure 2.**
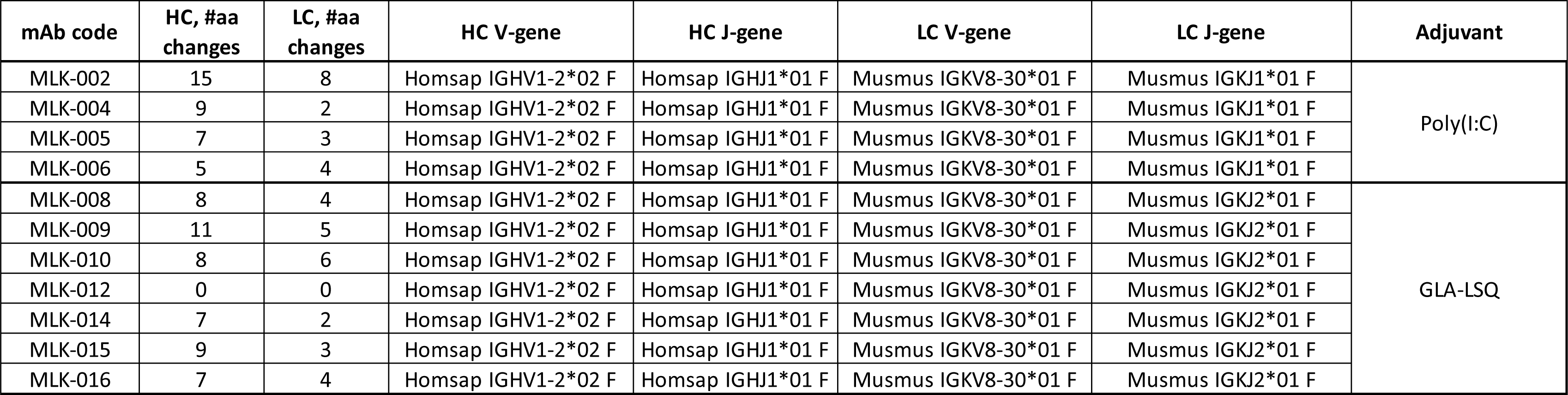
Information on the VRC01-like mAbs isolated following the boost immunization with HxB2.WT.Core. Eleven VRC01-like mAbs were generated from animals in the Poly(I:C) and GLA-LSQ groups following their immunization with the HxB2.WT.Core Ferritin. The Env-recognition properties of these mAbs is presented in Figure 5. Their neutralizing potentials are presented in **Table 1 B**.

**Supplemental Table 1.**
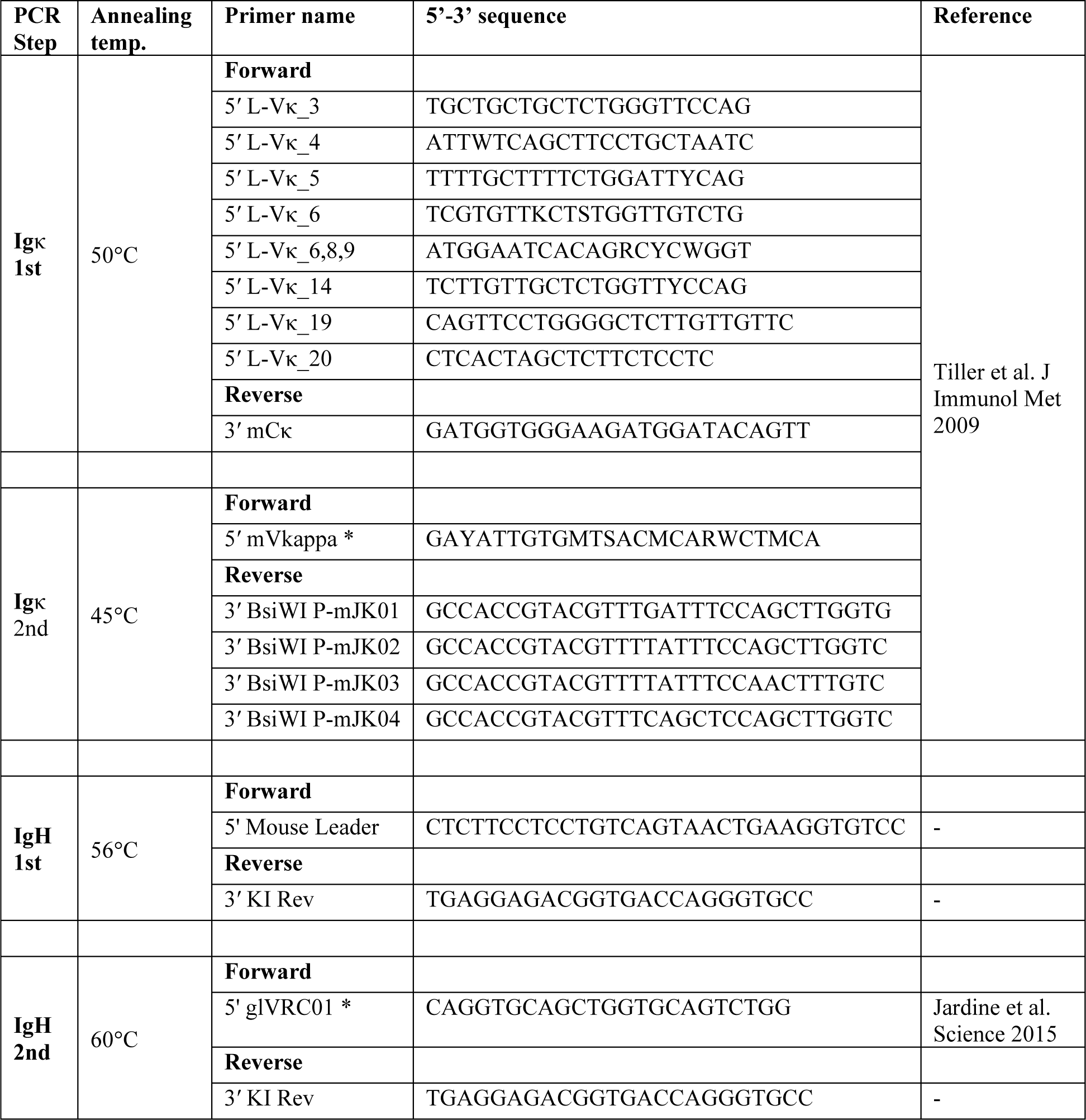

## REFERENCES

1. D. R. Burton, L. Hangartner, Broadly Neutralizing Antibodies to HIV and Their Role in Vaccine Design. Annu Rev Immunol 34, 635–659 (2016).

2. J. R. Mascola, B. F. Haynes, HIV-1 neutralizing antibodies: understanding nature’s pathways. Immunological reviews 254, 225–244 (2013).

3. A. P. West, Jr. et al., Structural insights on the role of antibodies in HIV-1 vaccine and therapy. Cell 156, 633–648 (2014).

4. J. N. Bhiman et al., Viral variants that initiate and drive maturation of V1V2-directed HIV-1 broadly neutralizing antibodies. Nat Med 21, 1332–1336 (2015).

5. N. A. Doria-Rose et al., Developmental pathway for potent V1V2-directed HIV-neutralizing antibodies. Nature 509, 55–62 (2014).

6. J. Gorman et al., Structures of HIV-1 Env V1V2 with broadly neutralizing antibodies reveal commonalities that enable vaccine design. Nat Struct Mol Biol 23, 81–90 (2016).

7. J. S. McLellan et al., Structure of HIV-1 gp120 V1/V2 domain with broadly neutralizing antibody PG9. Nature 480, 336–343 (2011).

8. M. Pancera et al., Crystal structure of PG16 and chimeric dissection with somatically related PG9: structure-function analysis of two quaternary-specific antibodies that effectively neutralize HIV-1. J Virol 84, 8098–8110 (2010).

9. L. M. Walker et al., Broad and potent neutralizing antibodies from an African donor reveal a new HIV-1 vaccine target. Science 326, 285–289 (2009).

10. H. B. Gristick et al., Natively glycosylated HIV-1 Env structure reveals new mode for antibody recognition of the CD4-binding site. Nat Struct Mol Biol 23, 906–915 (2016).

11. J. Huang et al., Identification of a CD4-Binding-Site Antibody to HIV that Evolved Near-Pan Neutralization Breadth. Immunity 45, 1108–1121 (2016).

12. P. D. Kwong, J. R. Mascola, Human antibodies that neutralize HIV-1: identification, structures, and B cell ontogenies. Immunity 37, 412–425 (2012).

13. M. M. Sajadi et al., Identification of Near-Pan-neutralizing Antibodies against HIV-1 by Deconvolution of Plasma Humoral Responses. Cell 173, 1783–1795 e1714 (2018).

14. J. F. Scheid et al., Sequence and structural convergence of broad and potent HIV antibodies that mimic CD4 binding. Science 333, 1633–1637 (2011).

15. X. Wu et al., Focused evolution of HIV-1 neutralizing antibodies revealed by structures and deep sequencing. Science 333, 1593–1602 (2011).

16. T. Zhou et al., Structural Repertoire of HIV-1-Neutralizing Antibodies Targeting the CD4 Supersite in 14 Donors. Cell 161, 1280–1292 (2015).

17. C. F. Barbas, 3rd et al., Recombinant human Fab fragments neutralize human type 1 immunodeficiency virus in vitro. Proc Natl Acad Sci U S A 89, 9339-9343 (1992).

18. H. J. Ditzel et al., Neutralizing recombinant human antibodies to a conformational V2- and CD4-binding site-sensitive epitope of HIV-1 gp120 isolated by using an epitope-masking procedure. J. Immunol. 154, 893–906 (1995).

19. J. Umotoy et al., Rapid and Focused Maturation of a VRC01-Class HIV Broadly Neutralizing Antibody Lineage Involves Both Binding and Accommodation of the N276-Glycan. Immunity 51, 141–154 e146 (2019).

20. C. Blattner et al., Structural delineation of a quaternary, cleavage-dependent epitope at the gp41-gp120 interface on intact HIV-1 Env trimers. Immunity 40, 669–680 (2014).

21. E. Falkowska et al., Broadly neutralizing HIV antibodies define a glycan-dependent epitope on the prefusion conformation of gp41 on cleaved envelope trimers. Immunity 40, 657–668 (2014).

22. J. Huang et al., Broad and potent neutralization of HIV-1 by a gp41-specific human antibody. Nature 491, 406–412 (2012).

23. L. Scharf et al., Antibody 8ANC195 reveals a site of broad vulnerability on the HIV-1 envelope spike. Cell reports 7, 785–795 (2014).

24. L. Scharf et al., Broadly Neutralizing Antibody 8ANC195 Recognizes Closed and Open States of HIV-1 Env. Cell 162, 1379–1390 (2015).

25. T. Schoofs et al., Broad and Potent Neutralizing Antibodies Recognize the Silent Face of the HIV Envelope. Immunity 50, 1513–1529 e1519 (2019).

26. T. Zhou et al., A Neutralizing Antibody Recognizing Primarily N-Linked Glycan Targets the Silent Face of the HIV Envelope. Immunity 48, 500–513 e506 (2018).

27. F. M. Brunel et al., Structure-function analysis of the epitope for 4E10, a broadly neutralizing human immunodeficiency virus type 1 antibody. J Virol 80, 1680–1687 (2006).

28. R. M. Cardoso et al., Broadly neutralizing anti-HIV antibody 4E10 recognizes a helical conformation of a highly conserved fusion-associated motif in gp41. Immunity 22, 163–173 (2005).

29. T. Muster et al., Cross-neutralizing activity against divergent human immunodeficiency virus type 1 isolates induced by the gp41 sequence ELDKWAS. J Virol 68, 4031–4034 (1994).

30. T. Muster et al., A conserved neutralizing epitope on gp41 of human immunodeficiency virus type 1. J Virol 67, 6642–6647. (1993).

31. G. Stiegler et al., A potent cross-clade neutralizing human monoclonal antibody against a novel epitope on gp41 of human immunodeficiency virus type 1. AIDS Res Hum Retroviruses 17, 1757–1765 (2001).

32. M. B. Zwick et al., Anti-human immunodeficiency virus type 1 (HIV-1) antibodies 2F5 and 4E10 require surprisingly few crucial residues in the membrane-proximal external region of glycoprotein gp41 to neutralize HIV-1. J Virol 79, 1252–1261 (2005).

33. M. B. Zwick et al., Broadly neutralizing antibodies targeted to the membrane-proximal external region of human immunodeficiency virus type 1 glycoprotein gp41. J Virol 75, 10892–10905 (2001).

34. J. H. Lee et al., Antibodies to a conformational epitope on gp41 neutralize HIV-1 by destabilizing the Env spike. Nature communications 6, 8167 (2015).

35. D. A. Calarese et al., Antibody domain exchange is an immunological solution to carbohydrate cluster recognition. Science 300, 2065–2071 (2003).

36. C. N. Scanlan et al., The broadly neutralizing anti-human immunodeficiency virus type 1 antibody 2G12 recognizes a cluster of alpha1-->2 mannose residues on the outer face of gp120. J Virol 76, 7306–7321 (2002).

37. A. Trkola et al., Human monoclonal antibody 2G12 defines a distinctive neutralization epitope on the gp120 glycoprotein of human immunodeficiency virus type 1. J Virol 70, 1100–1108. (1996).

38. M. Bonsignori et al., Staged induction of HIV-1 glycan-dependent broadly neutralizing antibodies. Science translational medicine 9, (2017).

39. K. J. Doores et al., Two classes of broadly neutralizing antibodies within a single lineage directed to the high-mannose patch of HIV envelope. J Virol 89, 1105–1118 (2015).

40. J. P. Julien et al., Broadly neutralizing antibody PGT121 allosterically modulates CD4 binding via recognition of the HIV-1 gp120 V3 base and multiple surrounding glycans. PLoS Pathog 9, e1003342 (2013).

41. H. Mouquet et al., Complex-type N-glycan recognition by potent broadly neutralizing HIV antibodies. Proc Natl Acad Sci U S A 109, E3268–3277 (2012).

42. M. Pancera et al., N332-Directed broadly neutralizing antibodies use diverse modes of HIV-1 recognition: inferences from heavy-light chain complementation of function. PLoS ONE 8, e55701 (2013).

43. R. Pejchal et al., A potent and broad neutralizing antibody recognizes and penetrates the HIV glycan shield. Science 334, 1097–1103 (2011).

44. L. M. Walker et al., Broad neutralization coverage of HIV by multiple highly potent antibodies. Nature 477, 466–470 (2011).

45. L. M. Walker et al., A Limited Number of Antibody Specificities Mediate Broad and Potent Serum Neutralization in Selected HIV-1 Infected Individuals. PLoS Pathog 6, 1–14 (2010).

46. L. M. Walker et al., Rapid development of glycan-specific, broad, and potent anti-HIV-1 gp120 neutralizing antibodies in an R5 SIV/HIV chimeric virus infected macaque. Proc Natl Acad Sci U S A 108, 20125–20129 (2011).

47. N. S. Longo et al., Multiple Antibody Lineages in One Donor Target the Glycan-V3 Supersite of the HIV-1 Envelope Glycoprotein and Display a Preference for Quaternary Binding. J Virol 90, 10574–10586 (2016).

48. P. L. Moore et al., Evolution of an HIV glycan-dependent broadly neutralizing antibody epitope through immune escape. Nat Med 18, 1688–1692 (2012).

49. L. Scharf et al., Structural basis for HIV-1 gp120 recognition by a germ-line version of a broadly neutralizing antibody. Proc Natl Acad Sci U S A 110, 6049–6054 (2013).

50. L. Scharf et al., Structural basis for germline antibody recognition of HIV-1 immunogens. eLife 5, (2016).

51. A. P. West, Jr., R. Diskin, M. C. Nussenzweig, P. J. Bjorkman, Structural basis for germ-line gene usage of a potent class of antibodies targeting the CD4-binding site of HIV-1 gp120. Proc Natl Acad Sci U S A 109, E2083–2090 (2012).

52. X. Wu et al., Maturation and Diversity of the VRC01-Antibody Lineage over 15 Years of Chronic HIV-1 Infection. Cell 161, 470–485 (2015).

53. T. Zhou et al., Multidonor Analysis Reveals Structural Elements, Genetic Determinants, and Maturation Pathway for HIV-1 Neutralization by VRC01-Class Antibodies. Immunity 39, 245-258 (2013).

54. M. D. Gray, et al., Characterization of a vaccine-elicited human antibody with sequence homology to VRC01-class antibodies that binds the C1C2 gp120 domain. bioRxiv, 2021.2008.2021.457217 (2021).

55. F. Klein et al., Somatic Mutations of the Immunoglobulin Framework Are Generally Required for Broad and Potent HIV-1 Neutralization. Cell 153, 126–138 (2013).

56. A. B. Balazs et al., Vectored immunoprophylaxis protects humanized mice from mucosal HIV transmission. Nat Med 20, 296–300 (2014).

57. M. Shingai et al., Passive transfer of modest titers of potent and broadly neutralizing anti-HIV monoclonal antibodies block SHIV infection in macaques. J Exp Med 211, 2061–2074 (2014).

58. L. Corey et al., Two Randomized Trials of Neutralizing Antibodies to Prevent HIV-1 Acquisition. N Engl J Med 384, 1003–1014 (2021).

59. S. Hoot et al., Recombinant HIV Envelope Proteins Fail to Engage Germline Versions of Anti-CD4bs bNAbs. PLoS Pathog 9, e1003106 (2013).

60. J. Jardine et al., Rational HIV immunogen design to target specific germline B cell receptors. Science 340, 711–716 (2013).

61. A. T. McGuire, J. A. Glenn, A. Lippy, L. Stamatatos, Diverse recombinant HIV-1 Envs fail to activate B cells expressing the germline B cell receptors of the broadly neutralizing anti-HIV-1 antibodies PG9 and 447-52D. J Virol 88, 2645–2657 (2014).

62. A. T. McGuire et al., Engineering HIV envelope protein to activate germline B cell receptors of broadly neutralizing anti-CD4 binding site antibodies. J Exp Med 210, 655–663 (2013).

63. L. Stamatatos, M. Pancera, A. T. McGuire, Germline-targeting immunogens. Immunological reviews 275, 203–216 (2017).

64. J. G. Jardine et al., HIV-1 broadly neutralizing antibody precursor B cells revealed by germline-targeting immunogen. Science 351, 1458–1463 (2016).

65. Y. R. Lin et al., HIV-1 VRC01 Germline-Targeting Immunogens Select Distinct Epitope-Specific B Cell Receptors. Immunity 53, 840–851 e846 (2020).

66. A. T. McGuire et al., Specifically modified Env immunogens activate B-cell precursors of broadly neutralizing HIV-1 antibodies in transgenic mice. Nature communications 7, 10618 (2016).

67. M. Medina-Ramirez et al., Design and crystal structure of a native-like HIV-1 envelope trimer that engages multiple broadly neutralizing antibody precursors in vivo. J Exp Med 214, 2573–2590 (2017).

68. K. R. Parks et al., Overcoming Steric Restrictions of VRC01 HIV-1 Neutralizing Antibodies through Immunization. Cell reports 29, 3060–3072 e3067 (2019).

69. A. T. McGuire et al., HIV antibodies. Antigen modification regulates competition of broad and narrow neutralizing HIV antibodies. Science 346, 1380–1383 (2014).

70. T. Zhou et al., Structural basis for broad and potent neutralization of HIV-1 by antibody VRC01. Science 329, 811–817 (2010).

71. A. J. Borst et al., Germline VRC01 antibody recognition of a modified clade C HIV-1 envelope trimer and a glycosylated HIV-1 gp120 core. eLife 7, (2018).

72. R. K. Abbott et al., Precursor Frequency and Affinity Determine B Cell Competitive Fitness in Germinal Centers, Tested with Germline-Targeting HIV Vaccine Immunogens. Immunity 48, 133–146 e136 (2018).

73. B. Briney et al., Tailored Immunogens Direct Affinity Maturation toward HIV Neutralizing Antibodies. Cell 166, 1459–1470 e1411 (2016).

74. X. Chen et al., Vaccination induces maturation in a mouse model of diverse unmutated VRC01-class precursors to HIV-neutralizing antibodies with >50% breadth. Immunity 54, 324–339 e328 (2021).

75. P. Dosenovic et al., Anti-HIV-1 B cell responses are dependent on B cell precursor frequency and antigen-binding affinity. Proc Natl Acad Sci U S A, (2018).

76. J. G. Jardine et al., HIV-1 VACCINES. Priming a broadly neutralizing antibody response to HIV-1 using a germline-targeting immunogen. Science 349, 156-161 (2015).

77. D. Sok et al., Priming HIV-1 broadly neutralizing antibody precursors in human Ig loci transgenic mice. Science 353, 1557–1560 (2016).

78. M. Tian et al., Induction of HIV Neutralizing Antibody Lineages in Mice with Diverse Precursor Repertoires. Cell 166, 1471–1484 e1418 (2016).

79. B. N. Lambrecht, M. Kool, M. A. Willart, H. Hammad, Mechanism of action of clinically approved adjuvants. Curr Opin Immunol 21, 23–29 (2009).

80. B. Pulendran, S. A. P, D. T. O’Hagan, Emerging concepts in the science of vaccine adjuvants. Nat Rev Drug Discov 20, 454–475 (2021).

81. M. Silva, et al., A particulate saponin/TLR agonist vaccine adjuvant alters lymph flow and modulates adaptive immunity. Sci Immunol 6, eabf1152 (2021).

82. C. Havenar-Daughton et al., The human naive B cell repertoire contains distinct subclasses for a germline-targeting HIV-1 vaccine immunogen. Science translational medicine 10, (2018).

83. C. C. LaBranche et al., HIV-1 envelope glycan modifications that permit neutralization by germline-reverted VRC01-class broadly neutralizing antibodies. PLoS Pathog 14, e1007431 (2018).

84. R. M. Lynch et al., HIV-1 fitness cost associated with escape from the VRC01 class of CD4 binding site neutralizing antibodies. J Virol 89, 4201–4213 (2015).

85. P. Dosenovic et al., Immunization for HIV-1 Broadly Neutralizing Antibodies in Human Ig Knockin Mice. Cell 161, 1505–1515 (2015).

86. D. S. Dimitrov, Therapeutic antibodies, vaccines and antibodyomes. mAbs 2, 347–356 (2010).

87. M. L. Salem, A. N. Kadima, D. J. Cole, W. E. Gillanders, Defining the antigen-specific T-cell response to vaccination and poly(I:C)/TLR3 signaling: evidence of enhanced primary and memory CD8 T-cell responses and antitumor immunity. J Immunother 28, 220–228 (2005).

88. L. Alexopoulou, A. C. Holt, R. Medzhitov, R. A. Flavell, Recognition of double-stranded RNA and activation of NF-kappaB by Toll-like receptor 3. Nature 413, 732–738 (2001).

89. R. N. Coler et al., Development and characterization of synthetic glucopyranosyl lipid adjuvant system as a vaccine adjuvant. PLoS One 6, e16333 (2011).

90. A. J. Radtke et al., Adjuvant and carrier protein-dependent T-cell priming promotes a robust antibody response against the Plasmodium falciparum Pfs25 vaccine candidate. Scientific reports 7, 40312 (2017).

91. S. G. Reed, D. Carter, C. Casper, M. S. Duthie, C. B. Fox, Correlates of GLA family adjuvants’ activities. Semin Immunol 39, 22–29 (2018).

92. A. L. Gavin et al., Adjuvant-enhanced antibody responses in the absence of toll-like receptor signaling. Science 314, 1936–1938 (2006).

93. S. L. Baldwin et al., Prophylactic efficacy against Mycobacterium tuberculosis using ID93 and lipid-based adjuvant formulations in the mouse model. PLoS One 16, e0247990 (2021).

94. T. Tiller et al., Efficient generation of monoclonal antibodies from single human B cells by single cell RT-PCR and expression vector cloning. J Immunol Methods 329, 112–124 (2008).

95. X. Brochet, M. P. Lefranc, V. Giudicelli, IMGT/V-QUEST: the highly customized and integrated system for IG and TR standardized V-J and V-D-J sequence analysis. Nucleic Acids Res 36, W503–508 (2008).

96. J. Snijder et al., An Antibody Targeting the Fusion Machinery Neutralizes Dual-Tropic Infection and Defines a Site of Vulnerability on Epstein-Barr Virus. Immunity 48, 799–811 e799 (2018).

97. C. C. LaBranche et al., Determinants of CD4 independence for a human immunodeficiency virus type 1 variant map outside regions required for coreceptor specificity. J Virol 73, 10310–10319. (1999).

